# The CoralAssist Plug: a novel device that enhances outplant rate and survivorship of sexually propagated corals

**DOI:** 10.64898/2026.08.27.747290

**Authors:** Eveline van der Steeg, Adriana Humanes, John C. Bythell, Alasdair J. Edwards, Yimnang Golbuu, Liam Lachs, Margaret W. Miller, James R. Guest

## Abstract

Sexual coral propagation is an emerging technique capable of producing large numbers of corals for coral transplantation and reef rehabilitation. In contrast to asexual coral propagation, sexual propagation increases genotypic diversity and can be used for selective breeding to enhance coral heat tolerance or other desirable traits. However, implementation at meaningful ecological scales is hindered by high mortality during early life stages, high costs associated with nursery rearing facilities, and labour-intensive outplanting methods. To overcome these issues, we developed the CoralAssist Plug (CAP), a ceramic device designed for the rapid and cost-effective outplanting of sexually propagated corals in large numbers that maximises post-outplant survivorship. CAPs combine three important functional features: 1) built-in microrefugia to protect juvenile corals from grazing, 2) a relatively small size, 3 by 1 cm, that is easy to handle and stack efficiently without compromising the survivorship of corals, and 3) a hole in the middle that facilitates handling and attachment. CAPs were settled with *Acropora* aff. *digitifera* and outplanted to a reef crest after 1 to 6 months of ex situ nursery rearing. A 3-person dive team was able to outplant ∼120 CAPs in one 90-minute shallow dive (just over 2 minutes per CAP per person). With longer nursery durations of 6 months, it was possible to achieve 36 % yield (i.e., the proportion of devices with a surviving coral) 4-years post-outplant. With nursery durations shortened to 1 month, we were able to attain 24 % yield 3-years post-outplant. Microrefugia significantly enhanced post-outplant survivorship leading to an 11 % increase in yield 4 years post outplant compared to devices without microrefugia. Outplanted corals that had reached adult size, were self-attached and were reproductively mature after 4 years. Our results suggest that CAPs can play a meaningful role in reef rehabilitation by efficiently introducing sexually propagated corals into natural populations with clear applications to assisted evolution techniques, such as selective breeding.

## Introduction

In the face of increasing coral loss due to climate change-related disturbances, coral reef rehabilitation is progressively being seen as an important management tool. Transplanting asexually produced coral fragments to rapidly increase coral cover and structural complexity is the most widely used technique to rehabilitate degraded reefs (Boström-Einarsson et al., 2020). While this approach is relatively straightforward to implement, asexual propagation may lead to restored populations of corals with high levels of clonality. Sexual coral propagation is a complementary approach with substantial potential, as it greatly increases genotypic diversity and offers several benefits compared to asexual fragmentation (Banaszak et al., 2023; Randall et al., 2020). For example, assisted evolution techniques such as selective breeding for adaptive traits can only be carried out via sexual reproduction (Drury et al., 2022; Humanes et al., 2024; Van Oppen et al., 2015). In addition, very large quantities of propagules can be collected during spawning events, providing large quantities of corals for transplantation while limiting impacts on source populations (de la Cruz & Harrison, 2017; Doropoulos et al., 2019). Although spawning timing predictions and husbandry techniques have improved over the last twenty years (Baird et al., 2021; Marhaver et al., 2023; Omori, 2019), the use of sexually reared corals for rehabilitation efforts is still largely experimental or at early trial stages of implementation. For sexual propagation to be successfully upscaled for reef rehabilitation efforts, there is a need to enhance the efficiency of outplanting and survivorship at early settled life stages (Banaszak et al., 2023; Vardi et al., 2021).

Coral sexual propagation involves either seeding competent larvae directly to reef substratum (de la Cruz & Harrison, 2017; Edwards et al., 2015; Heyward et al., 2002) or settling larvae onto appropriate substrates, i.e. seeding devices, for rearing and subsequent deployment to the reef. Seeding devices can either be deployed en masse and allowed to self-attach (Chamberland et al., 2017; Ramsby et al., 2025; Waters et al., 2025; Whitman, et al., 2025) or attached to the reef substratum by hand (Guest et al., 2014; Nakamura et al., 2011; Villanueva et al., 2012). The former method is widely seen as being more scalable due to the reduced labour required, however there is likely a trade-off between the method of deployment and early coral survivorship. Direct attachment methods, while more time consuming to deploy, allow more targeted deployment and will likely increase retention and survivorship. Over the past 2 decades, a range of direct attached settlement devices and outplanting techniques have been developed and trialled. Typically, these devices are secured to the reef with a hammer and nail, or a Coralclip^®^, or using epoxy inserted in a predrilled hole for those with cylindrical bases (Suggett et al., 2019). Examples of such devices include the ceramic grooved stackable ‘Coral Settlement Device’ (Okamoto et al., 2008), the ‘Coral Peg’ (Omori & Iwao, 2009) and ‘Coral plug-ins’ (Guest et al. 2014), which consist of a concrete settlement head mounted on a plastic tube or wall plug. Other approaches include attaching coral rubble to wall plugs using epoxy (Ligson et al., 2019; Villanueva et al., 2012), settling corals directly on wall plugs (Boch & Morse, 2012), and using 3D-printed ceramic devices designed to reduce fish grazing and algae overgrowth (Crawford et al., 2022).

Success of coral sexual propagation is typically measured as “yield” at a given time point after outplant (with yield being the proportion of devices outplanted that are retained and have at least 1 living coral). Device retention and coral survivorship are therefore the two main drivers of variation in yield. Mortality during the first few months post-settlement is often high in corals and has multiple causes, including predation and incidental grazing (Gallagher & Doropoulos, 2017; Penin et al., 2010), competition with benthic organisms (Tebben et al., 2014; Vermeij & Sandin, 2008), and sedimentation (Babcock & Smith, 2002). Fish grazing in particular has been found to substantially increase mortality of unprotected coral outplants (Miller & Hay, 1998; Rivas et al., 2021; van der Steeg et al., 2025). Mortality can be reduced by outplanting corals at an older age and larger size (Guest et al., 2014). However, extending the nursery period requires maintaining permanent land-based facilities or underwater structures, which increases labour and material costs (Schmidt-Roach et al., 2025). While caging juvenile corals can increase survivorship (e.g Baria et al., 2010; Nakamura et al., 2011); it can also lead to higher mortality due to increased algal competition (Steneck et al., 2014; Webster et al., 2015). Settlement devices with microrefugia which reduce vulnerability to grazing pressure have been shown to increase early survivorship (e.g. Doropoulos et al., 2016; Nozawa, 2008; Randall et al., 2021; Whitman et al., 2024).

Here we present and test the CoralAssist Plug (CAP), a novel settlement device designed to enable rapid outplanting of sexually propagated corals. CAPs incorporate microrefugia that promote larval settlement and protect young corals from grazing. Each CAP has a hole in the middle for efficient stacking on rods, facilitating their handling and maintenance during larval settlement, nursery rearing, and transport to outplant sites. The hole also allows rapid attachment to the reef without adhesive. Our aim was to design a settlement device to enhance settlement success, nursery capacity, outplant attachment efficiency, and post-outplant survivorship. To assess the effect of microrefugia on coral outplant survivorship, CAP performance was compared with that of a smooth control device without grooves. CAPs were settled ex situ with *Acropora* aff. *digitifera* larvae collected during spawning events in 2 consecutive years. Following ex situ nursery rearing, corals were outplanted at 1, 3, 4 and 6 months after settlement. Survivorship and growth were monitored on the reef for up to 4 years after spawning, and the scaled cost per surviving corals (aged 1 to 4 years) was estimated.

## Methods and Materials

### Design of the CoralAssist Plug

The CoralAssist Plug (CAP) is a device for settling, rearing and outplanting corals produced by in-vitro fertilisation (IVF). CAPs consist of a gear-shaped piece of steatite ceramic with a central hole produced by machine tooling. The CAPs are 30 mm in diameter, 10 mm in height, and the hole is 10 mm in diameter. The 10 teeth are 4 mm high by 5 mm wide, creating grooves of 4 mm deep, 4.5 mm wide on the outside and 2 mm on the inside (Figure 1 A & B). The central hole is designed for efficient stacking on rods during coral larval settlement and rearing in nurseries (e.g., flow-through aquaria, in situ nurseries, or larval pools) for a variable time before being outplanted to the reef (Figure 1 C). When CAPs are stacked on a rod, only the sides are available for larval settlement. This creates a total surface area of 15 cm^2^ available to larvae on the side of each CAP, of which 10 cm^2^ is inside the grooves. The grooves are designed to provide microrefugia where newly settled coral spat are protected from grazing. CAPs with settled corals are attached directly to the reef by inserting a cable tie masonry push mount (40.2 mm × 10 mm) through the central hole into a pre-drilled hole in the reef substratum and secured by hammering (Figure 1 E-G). To test the effect of the microrefugia on survivorship, an additional control device with the same dimension and from the same material but without grooves was designed. The control device had an exterior surface area of 9.4 cm^2^. Hereafter, the 2 types of devices are referred to as “CAP” for devices with microrefugia and “control” for smooth devices.

**Figure 1.**
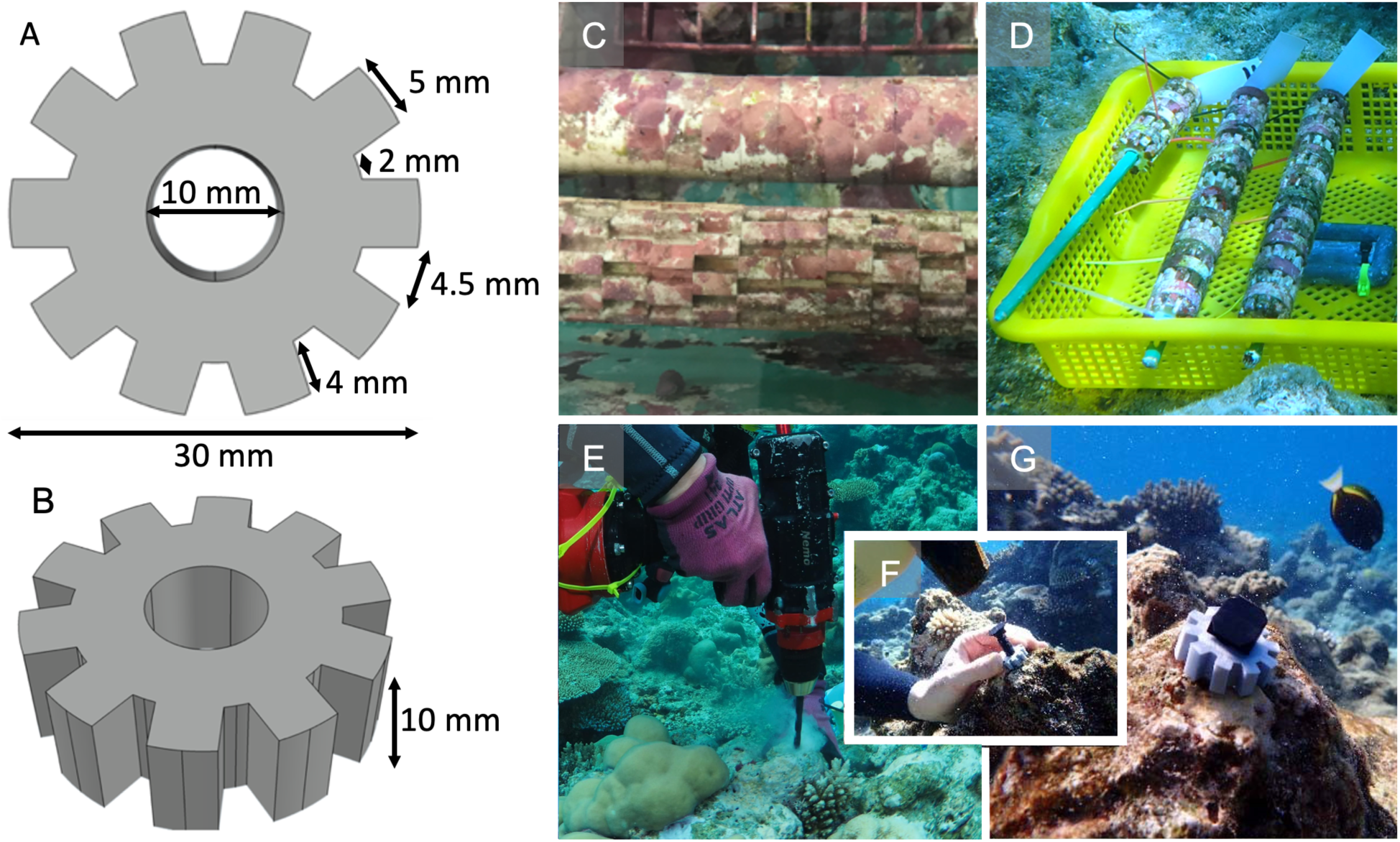
A & B) Design of the CoralAssist Plug (CAP), C) CAPs and controls in conditioning tank on rods, D) outplant baskets with CAPs and controls, E) outplanting of CAPs by drilling a hole, F) carefully tapping a push mount through the CAP to G) attach the CAP to the reef.

### Coral larval rearing

Spawning and rearing work was conducted at the Palau International Coral Reef Center (PICRC) in the Republic of Palau, West Pacific, in 2020 and 2021. Both coral collection and outplanting were carried out at Mascherchur, a shallow sheltered outer reef (N07°17ʹ29.3ʹʹ; E134°31ʹ8.00ʹʹ). The study species *Acropora* aff. *digitifera* (Dana, 1846) is a branching stony coral with high abundance on shallow exposed reefs in the Indo-West Pacific and known spawning timing in Palau (Baird et al., 2021; Gouezo et al., 2020; Penland et al., 2003).

In 2020, 10 *A.* aff. *digitifera* colonies containing visible pigmented oocytes with diameters over 20 cm were collected on April 3^rd^, 5 days before the full moon. The colonies were maintained in a 760 L shaded outdoor holding tank with a continuous flow of 50 μm filtered seawater (FSW). Water movement was provided by 2 magnetic pumps with each 3 flow accelerators (Pondmaster 1200 GPH, Accel Aquatic Vortex) and light by 4 LED aquarium lights on a 12:12 diurnal schedule (122 cm Reef Bright XHO 50/50). The colonies were checked nightly for spawning by turning the flow and pumps off from 19:00 till 21:30. Standard methods were used for gamete collection and larval rearing (Guest et al., 2010), for details see the Appendix (Appendix S1: Section S1). One hundred pre-conditioned devices (50 CAPs and 50 controls) were introduced to the larvae 4 days after spawning in seven 45 L tanks, whereas the 8^th^ tank had only 58 devices added due to limited availability (18 CAPs and 40 controls), thus 368 CAPs and 390 controls devices in total. The devices had been biologically conditioned for 6 months in two 184 L flow-through tanks with 2 pumps each (Hydor Koralia Nano 240 Circulation Pump/Powerhead). A mix of different species of crustose coralline algae (CCA) was collected from the study site and placed on top of the devices to encourage CCA coverage. The conditioning of devices with CCA in 1 tank was unsuccessful as the collected CCA died for unknown reasons, resulting in 2 different levels of conditioning: 1 set with a well-developed covering of CCA and another with little or no visible CCA (referred to subsequently as ‘biofilm conditioning‘). To avoid any bias during settlement, these 2 groups of devices were alternated on rods during settlement and the effect of conditioning on settlement success was tested (see description below).

After 1 week, no more swimming larvae were observed, and the devices were moved to 4 shallow 184 L ex situ nursery tanks until outplant after 3 or 6 months of nursery rearing (Figure 2A) for more details, see Appendix S1: Section S1. To compare the effects of CCA conditioning and device types on the number and survivorship of settled corals, a subset of 16 CAPs and 16 control devices, half with high CCA development and half with only biofilm conditioning, were assessed for coral settlement on day 11 and subsequent survivorship on days 25, 41 and 86 after settlement devices were introduced.

**Figure 2.**
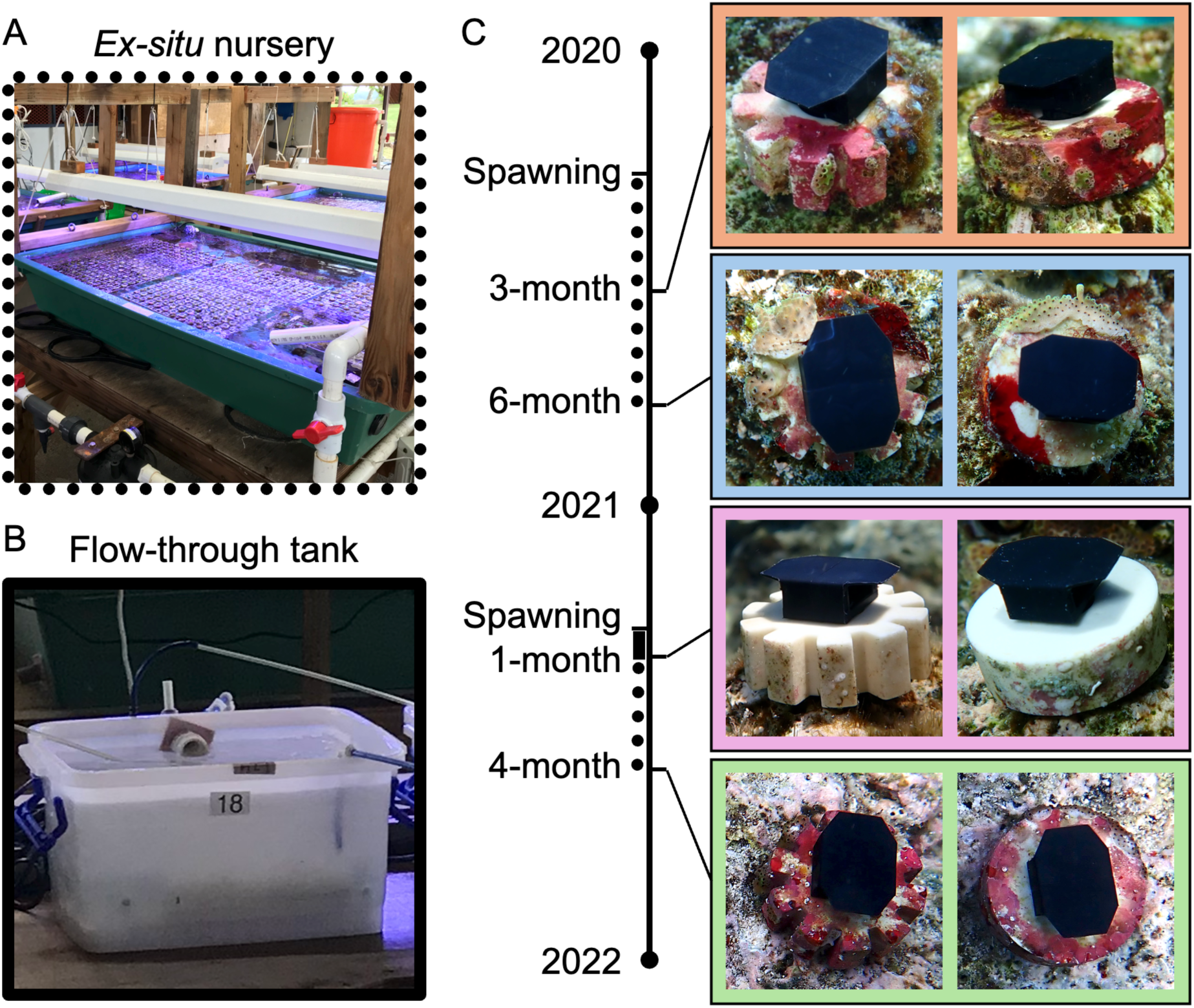
A) Ex situ nursery with submergible pumps, aquarium lights and micro herbivores, indicated on the timeline with dots and B) 10 L flow-through tank with aeration, indicated on the timeline with the solid bar. C) Timeline of coral spawning, nursery rearing and outplanting in this study with representative pictures of CoralAssist Plug (CAP) and control devices outplanted to the reef in the 4 outplants. The CAP and control devices have a diameter of 3 cm and a height of 1 cm. In 2020 devices were outplanted after a 3- and 6-month rearing period in ex situ nurseries and in 2021, the devices were kept in flow-through tanks until the first outplant after 1 month. The remaining devices were then moved to the ex situ nursery and outplanted after a 4-month rearing period.

In 2021, gravid *A.* aff. *digitifera* corals were collected on March 24^th^, 5 days before the full moon. On April 5th, 7 days after the full moon, 10 colonies spawned, and on the following night 1 colony spawned fully and 7 others had smaller releases. Coral larval rearing was carried out following the methods described above. Larvae were distributed among three 45 L settlement tanks to which 359 devices (171 CAPs and 186 control) with biofilm conditioning were distributed 3 days after spawning. At 1 week after settlement, the devices were moved to three 10 L tanks with constant 1 µm FSW flow through at 0.2 L/min and aeration for the first month (Figure 2 B). After 25 days, devices were assigned to 2 groups: the first group to be outplanted immediately with no further nursery rearing; and the second group to be outplanted after 4 months of rearing in the ex situ nursery tanks described above until they were outplanted. Parental colonies from the 2020 spawning were donated to the renewed Reef Crest display at the Palau Aquarium, whereas in 2021 they were secured to their original locations on the reef and nursery.

### Outplant and monitoring

Between 1 to 12 days prior to outplant, the number of living corals on each device was assessed and recorded using a stereo microscope. In 2020, devices were outplanted at the study site approximately 3 and 6 months (95-97 and 183-185 days) after spawning, with 200 devices at both timepoints (100 CAP and 100 control) (Figure 2 C). In 2021, devices were outplanted approximately 1 and 4 months (26 and 116 days) after spawning, with 150 devices (75 CAP and 75 control) and 148 devices (66 CAP and 82 control) per timepoint, respectively. For the devices outplanted at 3, 4 and 6 months, each had at least 1 living coral prior to outplant, whereas for the 1-month outplants, devices had at least 4 visible living corals to account for higher expected post-settlement mortality. To test if the phase in an ex situ nursery with lights and recirculation pumps could be eliminated, the 1-month-old CAPs in 2021 were outplanted straight from flow-through containers.

In all cases devices were outplanted on 10 m transects with a CAP and control plug alternating at 20 cm intervals. They were attached to the reef by drilling a hole into the bare reef substratum using a battery-powered submersible drill (Nemo Divers Drill) with an 8 mm drill bit (Figure 1 E). The substratum was brushed to remove the sediment from drilling and any turf algae. Nylon cable tie masonry push mounts (40.2 × 10 mm, Screwfix UK) were pushed through the devices before inserting the end of the mount into the hole. The mounts were carefully tapped with a hammer to ensure a tight fit of the devices to the reef (Figure 1 F & G).

The outplants from 2020 were surveyed for retention, status (dead, alive or missing), the number of live corals and self-attachment to the substratum. An outplant was considered dead when there were no live corals on the device. Surveys were conducted at 184 (first outplant only), 283, 394, 488, 565, 666, 741, 1089 and 1513 days after spawning and those from 2021 at 124 (first outplant only), 204, 301, 378, 725 and 1149 days after spawning. In 2022, 2023 and 2024, the maximum perpendicular diameters were measured using Vernier calipers for each living coral. The geometric mean diameter (GMD) of each coral was calculated by the formula:

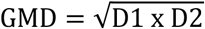

where, D1 is the maximum diameter, and D2 is the maximum perpendicular diameter (Guest et al., 2014). When multiple corals were present on a device, the GMD of the largest coral was included in the analysis. Photos were taken of each device during every survey as reference material in case the order on the reef was unclear if devices went missing. Self-attachment was defined as a coral growing from the device onto the surrounding reef substratum. Opportunistic gravidity checks were performed on 6 of the largest corals in 1 transect in 2024. There were 13 observations of newly recruited single-polyp coral spat on devices, which were excluded from the analysis.

### Data analysis

A generalized linear mixed-effects model (GLMM, Brooks et al. 2017) with a negative binomial distribution was used to test the effect of conditioning level (CCA vs. biofilm) on corals per device in the 2020 subset. Days after settlement (continuous), device type (2 levels), and conditioning type (2 levels) were included as fixed effects, with settlement tank (8 levels) as a random effect. Coral numbers at outplant were log-transformed and compared between device types and rearing duration using a Kruskal-Wallis test followed by a Bonferroni-corrected pairwise Wilcoxon rank-sum test. The effect of the log-transformed number of corals per device at outplant (continuous) and either device type or CAP outplant age (4 levels) on the number of corals per device at 1-year-old was tested using a Poisson GLMM (Bates et al., 2015) with transect (14 levels) as a random effect. The models were assessed for overdispersion and ability to predict zeros.

Survivorship between devices and outplant ages was compared using Kaplan-Meier survival analysis with right-censored data (Lee & Wang, 2003). When mortality occurred between surveys, the day of death was estimated as the midpoint between surveys. Differences between survivorship curves were tested using pairwise log-rank comparisons. Device retention, yield (retained devices containing ≥ 1 live coral, Chamberland et al. (2017)) and self-attachment at the end of the study were analyzed separately for each outplant year (2020, 2021) using binomial GLMMs with a logit link (Bates et al., 2015). Device type (2 levels) and outplant age (2 levels) were included as fixed effects and transect as a random intercept (8 levels in 2020; 6 levels in 2021). Although all 2021 models and the 2020 retention model produced near-zero transect variance (singular fits), the random effect was retained for consistency. Minor deviations in residual uniformity or variance in the 2020 retention and 2021 yield models were considered acceptable for binomial GLMMs, where groupwise differences in residual spread are expected and did not indicate problems with model fit. Annual detachment rates were compared descriptively as percentage detachment per transect standardized to yearly rates.

The effect of device type and outplant age on the GMD of the largest live coral per device at the end of the study (4 and 3 years, for 2020 and 2021 outplants, respectively) was tested using a Gaussian GLMM with a log link function and transect as a random effect (Brooks et al., 2017). A second model evaluated GMD at 3-years-old across all outplants, including device type (2 levels) and outplant age (2 levels) as fixed effects and transect and spawning year as random effects. Pairwise comparisons among treatments were performed using post hoc Tukey Tests (Lenth, 2017). All model validation steps included assessing residual homogeneity, dispersion, zero inflation using simulation-based diagnostics (Hartig, 2018). Analyses were performed in R using Rstudio (Rstudio Team 2010; R Core Team 2022). Values are reported as mean ± standard error unless otherwise indicated.

### Cost analysis – experimental and realistic rehabilitation costs

The experimental cost estimations for this study were carried out following Edwards (2010) and Humanes et al. (2021). The costs of capital equipment, consumables, labour, boat and SCUBA tank rental were separated. All capital equipment costs were divided by three, as most projects are long-term, and it was assumed all equipment would be used for at least 3 years. The wages were based on the rates in Palau at the time the study was carried out (2020-2021). This study was done over 2 years, so for a more straightforward cost analysis, we calculated the cost for 900 devices to be settled, reared and outplanted in 1 year rather than 2 years. Here we used a scenario where 200 devices were outplanted at 1-, 3- and 6-months-old as this was realistic with nursery yields and equivalent to the outplant ages in this study. The 1-month-old outplant was costed to be reared in basic flow-through tanks, and the 3- and 6-month-old outplants were costed to be reared in nurseries with aquarium lights and recirculation pumps. Monitoring was costed at yearly intervals. Cost estimations were separated by device type as survivorship differed between the CAP and control devices. The cost per live coral for the different nursery durations was estimated in the nurseries before outplant, at outplant, and yearly on the reef for each scenario.

Realistic rehabilitation cost was estimated based on a coral rehabilitation effort where 10,000 CAPs would be produced, settled, and outplanted. At this larger volume, the cost per CAP changed from $2.18 plus tooling costs to $1.01 using a cordierite-mullite ceramic. Only the realistic rehabilitation costs are presented in the main text of this study as small scale experimental costs are not representative of actual rehabilitation costs due to economies of scale (Humanes et al., 2021; Suggett et al., 2024). Realistic rehabilitation costs were calculated for three outplant scenarios as if all CAPs were outplanted at 1-, 3- or 6-months-old. CAPs in the 1-month-old outplant scenario were maintained in basic flow through containers until outplant, whereas the CAPs outplanted at 3- and 6-months-old were maintained in large holding tanks with aquarium lights and recirculation pumps. The outplanting was costed for 2 dive teams of 3 divers outplanting a total of 700 devices per day during three 90-minute dives. The number of devices outplanted varied for the 3 different nursery rearing times, as nursery yield reduced over time. Hence there were fewer devices to outplant for longer nursery durations. The monitoring cost was calculated for a subset of 400 CAPs to be monitored yearly, as it would not be feasible to monitor all outplanted devices in a large-scale outplant. For the rehabilitation outplant, only the cost for CAPs was estimated as they had higher survivorship in all outplants compared to the control devices. Details of the cost analysis are supplied in Appendix S2.

## Results

### Coral rearing

In 2020, an estimated total of 872,500 eggs were released by 3 colonies and fertilisation success of 94.6 ± 0.8 % was achieved (values are mean ± SE unless otherwise specified). Larval survivorship was 77.0 ± 7.9 % at 4 days after fertilisation. At 11 days after devices were introduced to larvae, devices with CCA conditioning had an average of 50.0 ± 15.2 settled coral spat per CAP and 29.6 ± 11.0 per control device. In contrast, devices with only biofilm conditioning had lower settlement with an average of 5.1 ± 2.7 spat per CAP and 3.5 ± 1.7 spat per control. When combined, this gives an estimated settlement success of 44.1 ± 11.4 %. Of all devices assessed in the subset, 84.4 % had at least 1 living coral spat (87.5 % and 81.3 % for CAPs and controls respectively). In 2021, 6 colonies released an estimated 734,000 eggs, and 96.0 ± 2.4 % fertilisation success was achieved. At 4 days after fertilisation, larval survivorship was 65.3 ± 3.8 %. Between 11 to 20 days after settlement, CAPs had an average of 10.6 ± 1.5 and control 11.7 ± 1.7 spat per device, giving a combined larval settlement success of 22.4 ± 2.3 %. A total of 86.0 % of devices had at least 1 living coral after settlement (85.5 % and 86.0 % for CAPs and controls respectively).

### Relationship between device and conditioning type on the number of corals per device over time

Both device type (CAP or control) and conditioning significantly influenced the mean initial number of corals per device (GLMM, CAP vs control: p = 0.006, estimate = 0.7, SE = 0.26, z-ratio = 2.74, biofilm vs CCA: p < 0.001, estimate = −1.96, SE = 0.28, z-ratio = −1.11, Figure 3, Appendix S1: Table S1). Devices with CCA conditioning had significantly more coral spat 11 days after settlement, with an average of 50.0 ± 15.2 and 29.6 ± 11.0 spat for CAPs and control devices respectively. For devices with biofilm this was 5.1 ± 2.7 and 3.5 ± 1.7 respectively (Figure 3). The half-life (50 % reduction in corals per device) was 55 and 58 days for CCA and biofilm conditioning respectively. At 3 months after settlement, CAPs had 68 % more corals per device for the CCA and 40 % for the biofilm conditioning compared to controls.

**Figure 3.**
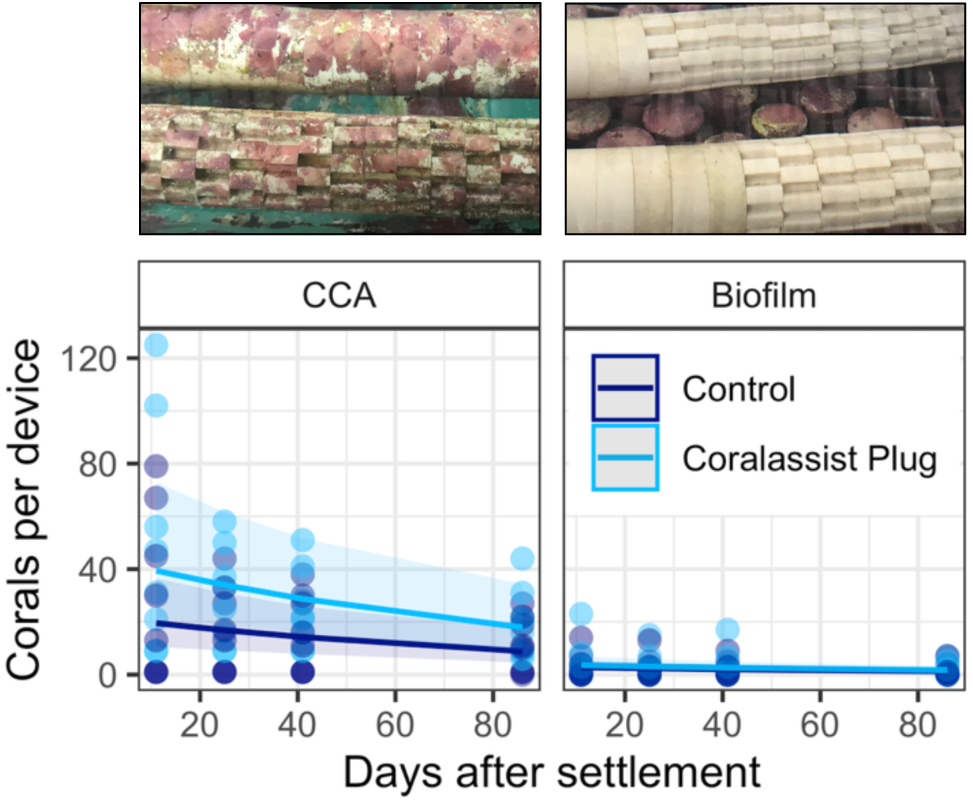
The expected relationship between the number of corals per device and days after settlement per device and conditioning type derived from a negative binomial GLMM. Shaded areas indicate the 95% confidence intervals; raw values are depicted as points. n = 8 devices per treatment and conditioning type, n = 32 total. Conditioning type for each model is depicted above the respective plot.

### Effect of coral density on CAPs and nursery duration on the number of corals after 1 year

The number of corals per device at outplant did not differ between device types (Kruskal–Wallis, χ^2^ = 0.044, df = 1, p = 0.834) but was significantly different between outplant ages (Kruskal–Wallis, χ^2^ = 232.3, df = 3, p < 0.001). Pairwise comparisons indicated that all outplants differed significantly (Wilcoxon rank sum test with Bonferroni correction, all p < 0.001) except between the 4- and 6-month-old outplants (p = 1.000, Appendix S1: Figure S1 and Table S2). The number of corals per CAP at the time of outplant positively affected the number of corals at 1 year old (GLMM, p < 0.001, estimate = exp (0.52) ≈ 1.68, SE = 0.06, z-ratio = 8.3; Figure 4, Appendix S1: Table S3). When comparing CAPs from different outplants, there was a significant difference in this relationship between the 3- and 6-month-old outplants from 2020 (GLMM, p < 0.001, estimate = exp (-0.63) ≈ 0.53, SE = 0.16, z-ratio = -4.31), the 1- and 4-month-old outplants from 2021 (GLMM, p = 0.013, estimate = exp (-0.65) ≈ 0.53, SE = 0.21, z-ratio = -3.02) and the 6- and 1-month-old outplants (GLMM, p < 0.001, estimate = exp (1.03) ≈ 2.80, SE = 0.19, z-ratio = 5.35) (Figure 4, Appendix S1: Table S3 and S4). Devices with a longer nursery time needed fewer corals at outplant to have at least 1 coral alive at 1 year old. CAPs outplanted after a 6- and 3-month nursery period in 2020 needed an average of 2 and 8 corals respectively. In 2021, the CAPs outplanted after 4- and 1-month rearing period needed 5 and 15 corals respectively (Figure 4). When the data was grouped per device type, the CAPs needed fewer corals than the control devices at outplant to obtain at least 1 live coral per device after 1 year (GLMM, p < 0.001, estimate = exp (0.32) ≈ 1.38, SE = 0.09, z-ratio = 3.77; Appendix S1: Figure S2 and Table S3). Apparent increases in the number of corals per device were due to partial mortality (e.g. fish grazing), where a single coral to split into two corals that were then counted separately (Figure 4).

**Figure 4.**
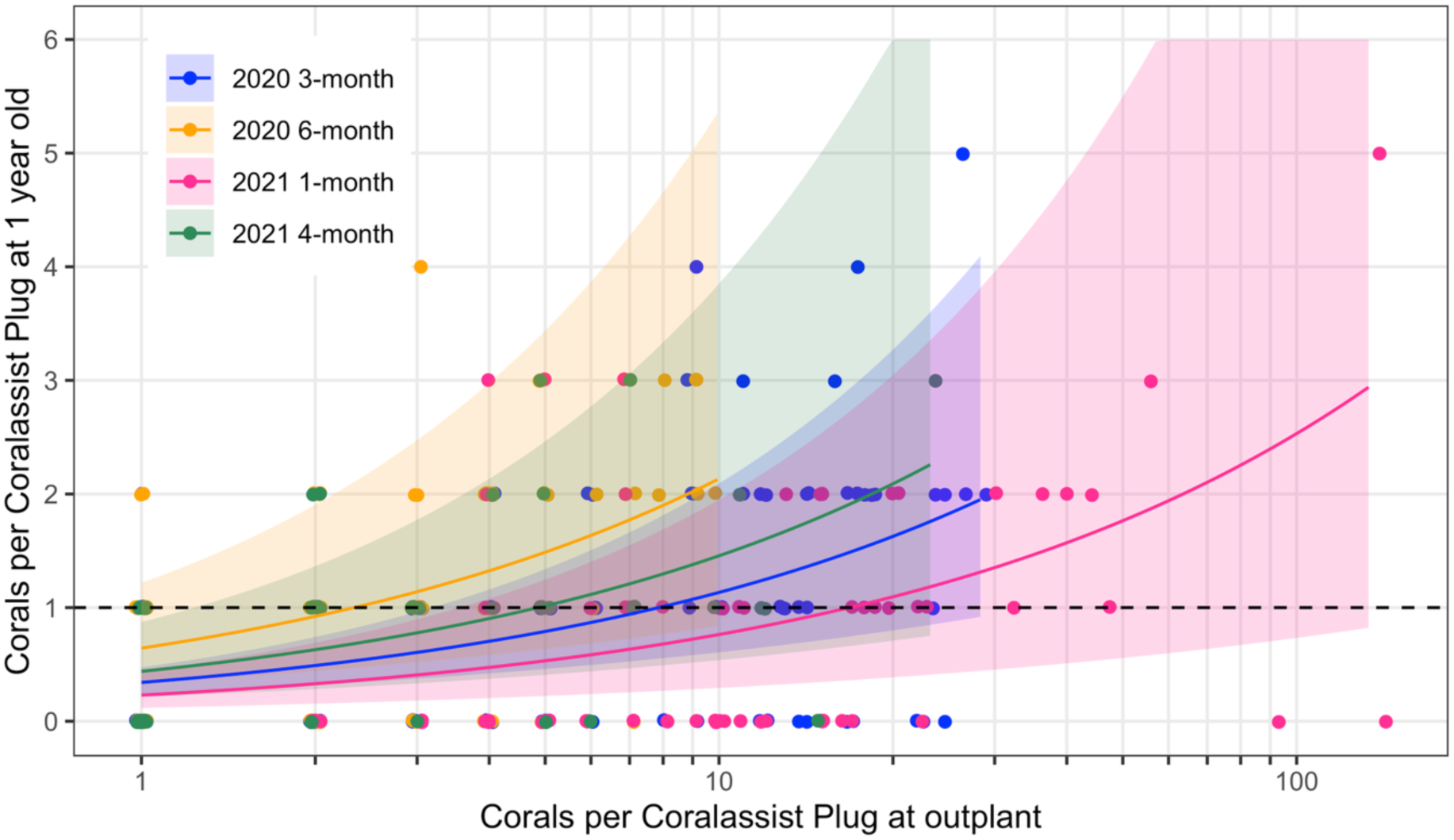
The relationship between corals per CoralAssist Plug (CAP) at outplant and at 1 year old, grouped per outplant age, derived from a GLMM with a Poisson distribution. The original data is represented with a jitter to prevent overlap, and the shaded areas represent 95 % confidence intervals. The dashed line represents 1 coral alive at 1 year old.

### Outplant time

With 2 divers, it took 164 ± 6 diver-seconds to outplant 1 device; with 3 divers, the deployment time was decreased to 138 ± 10 diver-seconds per device. This resulted in a deployment rate of 66 or 117 devices outplanted per 90-minute dive, or 33 or 39 devices per diver per dive, with 2 or 3 divers respectively.

### Survivorship, retention, yield and coral self-attachment

CAPs from the 6-month-old outplant had the highest mean survival time at 1049 ± 58 days SE (Figure 5 A, Appendix S1: Figure S3 displays the survival curves with confidence intervals, Appendix S1: Table S5). Their survival was significantly higher compared to the 3-month-old CAP and control outplants from 2020 (798 ± 59 days, p = 0.044 and 635 ± 58 days, p < 0.001, respectively), the 1-month-old CAP and control outplants from 2021 (546 ± 49 days, p < 0.001 and 483 ± 50 days, p < 0.001, respectively) and the 4-month-old control outplants from 2021 (610 ± 49 days, p = 0.002, Appendix S1: Table S5 and S6). The 6-month-old control outplants from 2020 and 4-month-old CAP outplants from 2021 (810 ± 61 days and 694 ± 57 days, respectively) also had higher survivorship compared to the 1-month-old control outplants from 2021 (483 ± 50 days, p = 0.033 and 0.018, respectively, Appendix S1: Table S5 and S6). When the 4 outplants were combined by device type, CAPs had significantly higher survivorship than the control devices (845 ± 33 days and 681 ± 31 days, p < 0.001, Appendix S1: Figure S4 and Table S7). CAPs outplanted at 1-, 3- 4- and 6-months-old had 25, 34, 49 and 61 % survivorship respectively after 3 years, and CAPs outplanted at 3-and 6-month-old had 32 % and 52 % survivorship after 4 years, compared to 19, 28, 32, 40 % and 24 and 40 % for control devices respectively.

**Figure 5.**
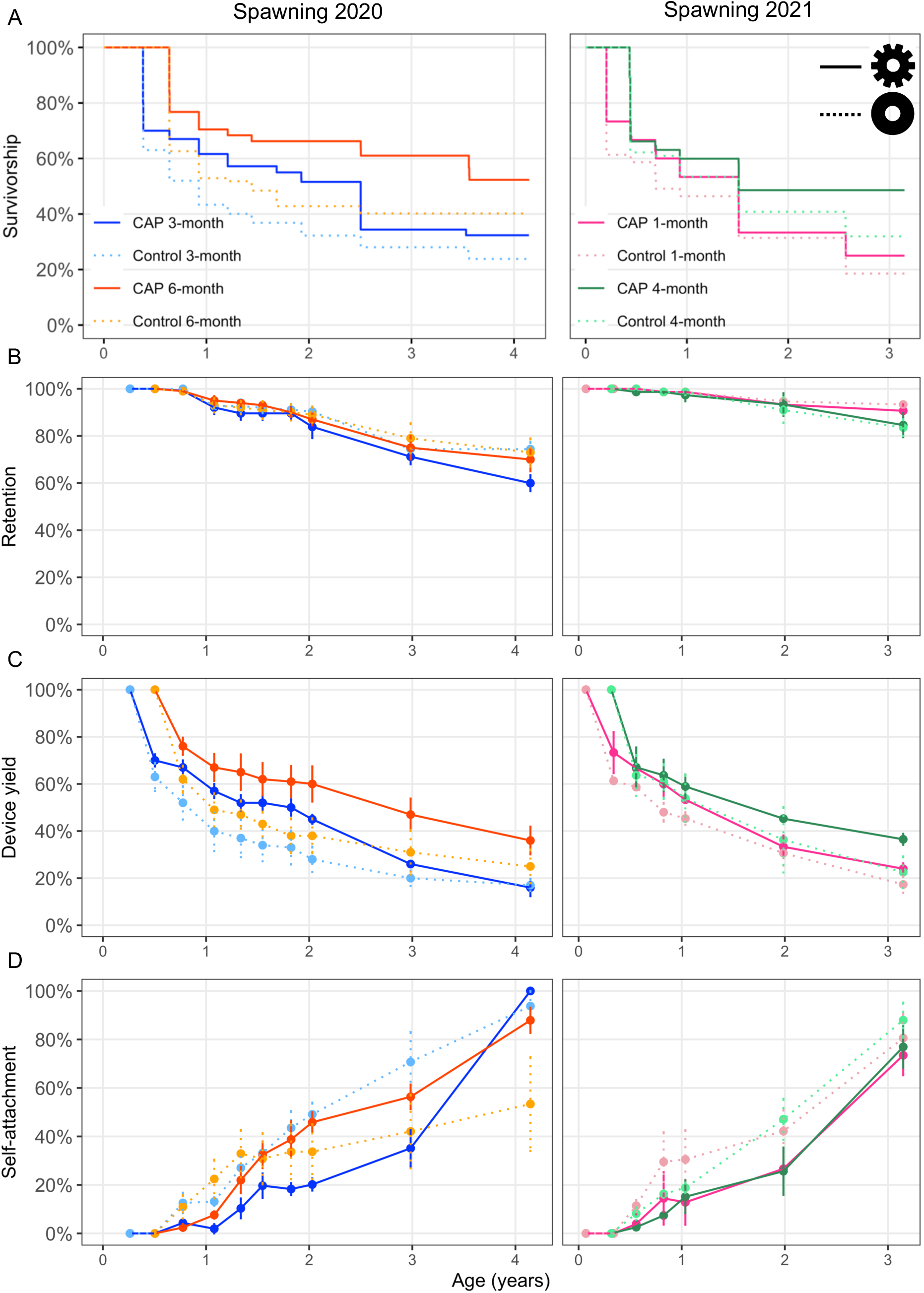
A) Device level survivorship curves of CoralAssist Plugs (CAP) and control devices outplanted to the reef after varying nursery periods in 2020 and 2021. Day = 0 is the day of spawning. Solid lines are the CAPs, and dashed lines are the control devices. B) Retention – the percentage of devices found throughout the study. C) Device yield – the percentage of outplanted devices that could be found and had ≥ 1 live coral. D) The percentage of devices with live corals that self-attached to the reef by growing onto the reef substratum. Error bars in B-D are the standard error, calculated for the different transects.

There was no effect of device type on retention in both outplant years, and between outplants in 2020. In the 2021 outplant, 1-month-old outplants had higher retention than the 4-month outplant (GLMM, p = 0.031, estimate = -0.81, SE = 0.38, z-ratio = -2.16, Figure 5 B, Appendix S1: Table S8 and S9). Taken over the entire study, the average yearly device detachment rate was 7.0 ± 0.9 % for CAPs and 5.6 ± 0.7 % for control devices. In the 2020 outplant, there was no difference in device yield between device types, but the 6-month-old outplant had a higher yield than the 3-month-old outplant (GLMM, p = 0.010, estimate = 0.80, SE = 0.31, z-ratio = 2.56, Figure 5 C, Appendix S1: Table S10). In the 2021 outplant, CAPs had a higher device yield compared to control devices (GLMM, p = 0.036, estimate = 0.57, SE = 0.27, z-ratio = 2.10, Appendix S1: Table S11), but there was no difference between outplant ages. At the end of the study, the percentage of self-attached corals was similar between the device types in both outplant years. In 2020 devices outplanted at 3-months-old had a higher self-attachment rate than those outplanted at 6-months-old (GLMM, 6-vs 3-month outplant: p = 0.031, estimate = −2.35, SE = 1.09, z-ratio = −2.16, Figure 5D, Appendix S1: Table S12 and S13), and there was no difference in self-attachment rate between outplant ages in 2021.

### Coral size

Neither device type nor rearing duration significantly influenced the GMD of the largest coral per device on corals from 2020 at 4-years-old (GLMM, CAP vs Control: p = 0.829, estimate = 0.01, SE = 0.06, z-ratio = 0.22, 3-vs 6-month outplant: p = 0.292, estimate = −0.12, SE = 0.10, z-ratio = −1.06, Appendix S1: Table S14) nor on corals from 2021 at 3-years-old (GLMM, CAP vs Control: p = 0.383, estimate = -0.06, SE = 0.07, z-ratio = -0.87, 1-vs 4-month outplant: p = 0.170, estimate = 0.16, SE = 0.12, z-ratio = 1.37, Appendix S1: Table S15)(Figure 6 A). When comparing all outplants at 3-years-old, the CAPs outplanted at 4-months-old had a larger GMD than CAPs outplanted at 3- and 6-months, and control devices outplanted at 4-months-old had a larger GMD than CAPs outplanted at 3-months-old and control devices outplanted at 3- and 6-months-old (Tukey test, p = 0.009, 0.022, 0.017, 0.009, 0.022, respectively, Appendix S1: Table S16). The mean GMD of corals on devices outplanted at 3- and 6-months-old was 4.9 ± 0.3 and 5.1 ± 0.2 cm after 3 years, and 10.3 ± 0.6 and 9.3 ± 0.3 cm after 4 years respectively, and the mean GMD of corals outplanted at 1- and 4-months-old was 5.6 ± 0.3 and 6.6 ± 0.3 cm after 3 years, respectively (Figure 6 B-E). The largest 3-year-old corals from the 1-, 3-, 4- and 6-month outplants reached a GMD of 10.3, 9.4, 12.8 and 8.5 cm respectively, and the largest 4-year-old corals from the 3- and 6-month outplants 20.4 and 14.6 cm, respectively (Figure 6 F-I). While a full survey of reproductive status was not possible, opportunistic checks found pigmented eggs in three out of six 4-year-old colonies in 2024.

**Figure 6.**
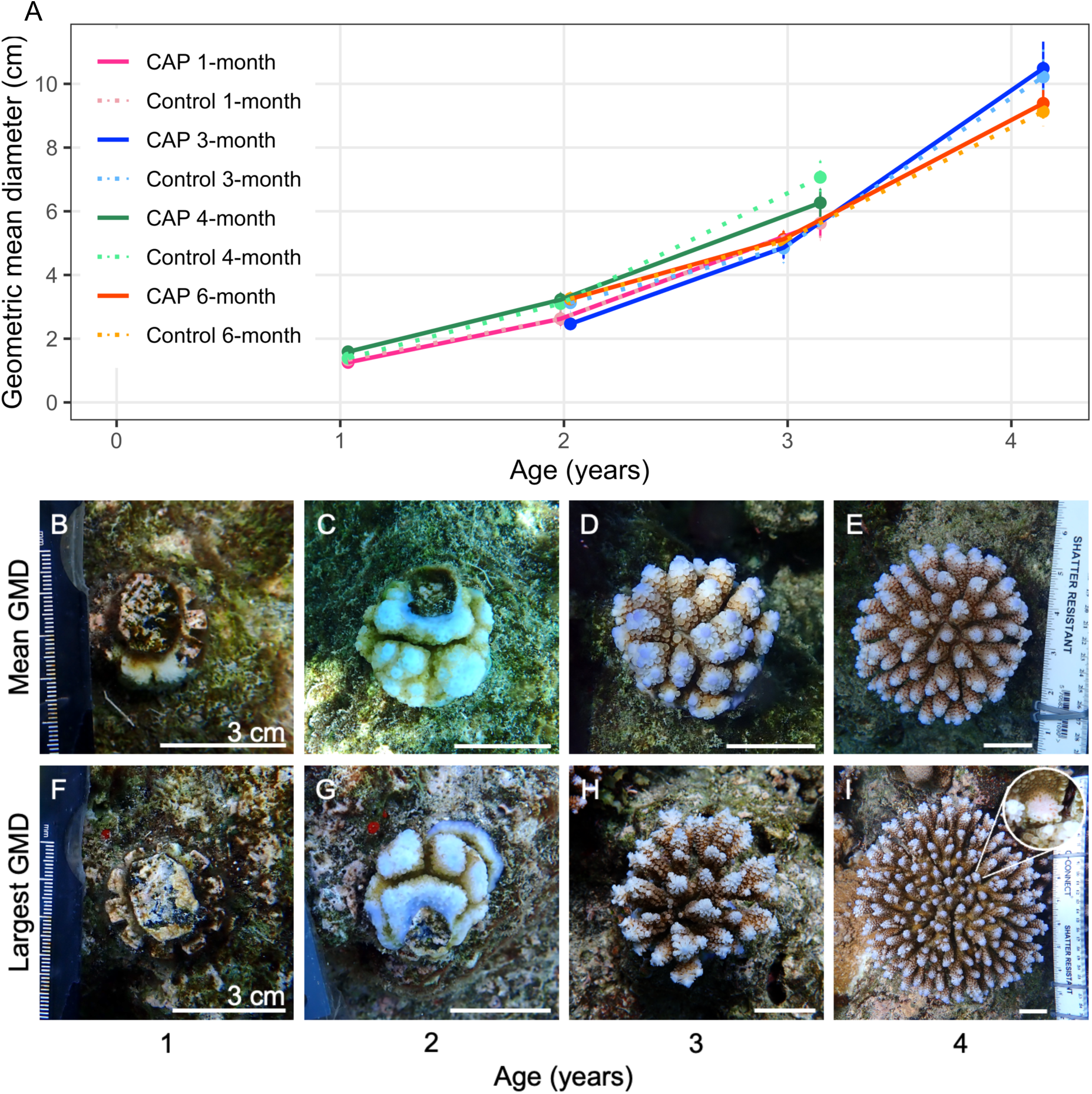
A) Change in geometric mean diameter (GMD) of the largest live coral per device in cm throughout the study (mean ± standard error). B-E) Time series of a CAP outplanted at 6-months-old with a GMD of 10.6 cm at 4 years old, close to the mean GMD. F-I) Time series of a CoralAssist Plug (CAP) outplanted at 3 months old with the largest GMD of 20.4 cm at 4 years old. I) After 4 years this coral was mature with pigmented eggs.

### Realistic rehabilitation cost estimates

The realistic rehabilitation production cost of a CAP with a live coral for the 1-month flow through tanks and 3- and 6-month ex situ nurseries was calculated at different stages (Table 1, Appendix S2). The total rehabilitation production costs for settling 10,000 CAPs were estimated to be $37,456, $41,520 and $41,154 after 4 years for the 1-, 3- and 6-month-old outplants, respectively. Total production costs were slightly lower for 6-month-old outplants than for 3-month-old outplants (Appendix S2). The higher nursery rearing costs of the additional 3 months were offset by the need for 2 fewer outplanting days, as mortality between 3 and 6 months reduced the nursery yield and therefore the number of CAPs to be outplanted. Additionally, the high yield of the CAPs with 6 months of nursery rearing at all timepoints (Figure 5 C) resulted in the lowest costs per coral from 2 years old onwards.

**Table 1.**
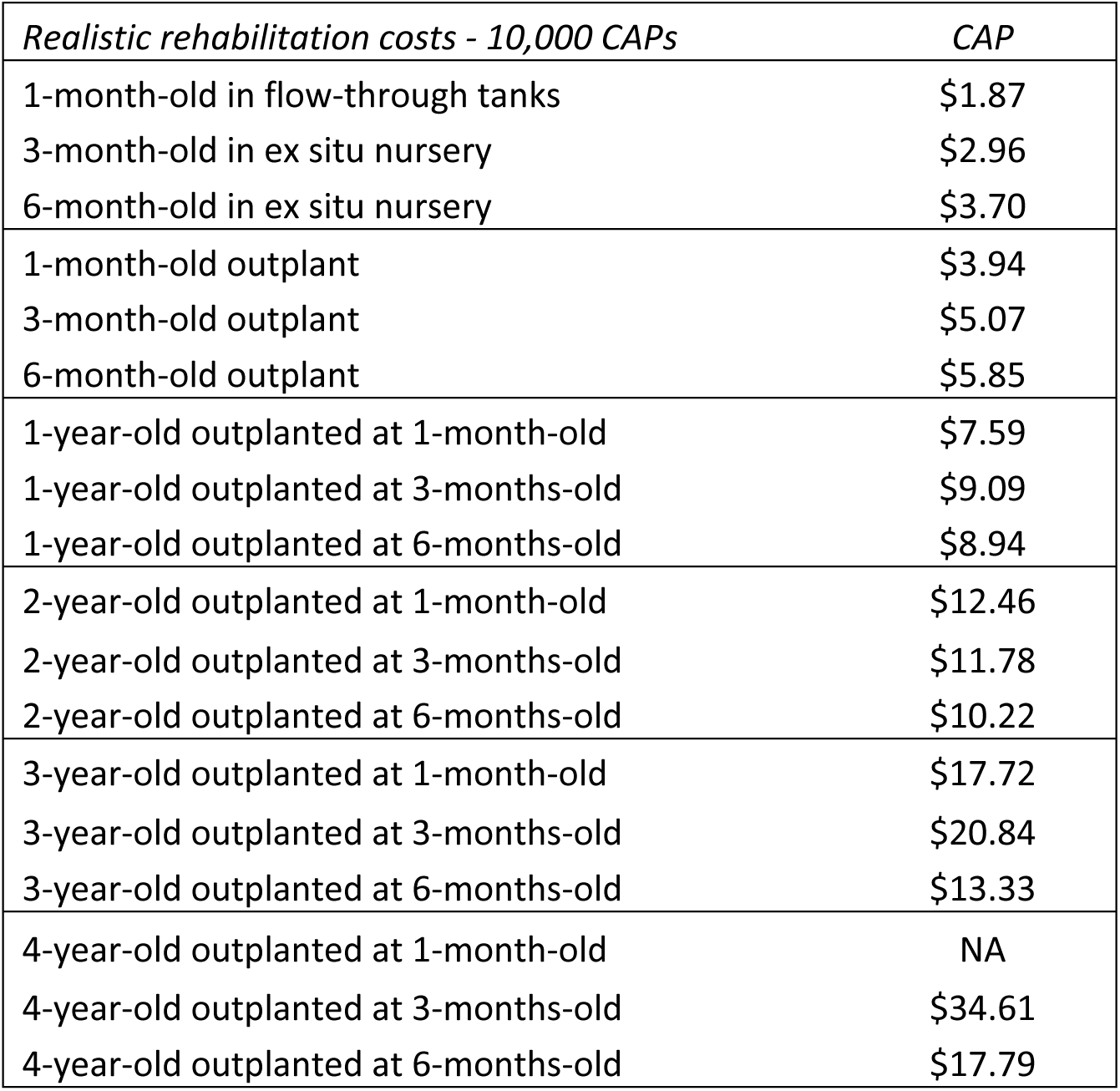
Estimated cost per CAP with at least 1 live coral reared with different nursery types and durations for a realistic rehabilitation initiative. Costs were estimated at different production stages and ages after being outplanted to the reef.

## Discussion

Sexual coral propagation is increasingly seen as one of the most promising approaches for producing coral outplants as part of reef rehabilitation efforts. Substantial progress has been made in refining protocols for predicting spawning (Baird et al., 2021; Mera et al., 2025), controlling spawning times (Craggs et al., 2017; Koukoumaftsis et al., 2025), ex situ fertilisation (Guest et al., 2010), larval husbandry (Pollock et al., 2017), settlement (Abdul Wahab et al., 2023) and outplant techniques (Mendoza Quiroz et al., 2025). Nonetheless, high post-outplant mortality and low retention of outplanted devices are major bottlenecks to scalability (Banaszak et al., 2023; Randall et al., 2020). Here we develop and demonstrate a novel device for coral sexual propagation - the CoralAssist Plug (CAP) - designed for ease of handling, rapid deployment and reduced post-outplant mortality due to fish grazing. We show that a small team of 3 divers could outplant several hundred CAPs per day. Furthermore, incorporating microrefugia into the device significantly improved survivorship, highlighting the importance of protection from grazers during early life stages. Extending the ex situ nursery rearing period (up to 6 months) further increased overall survivorship. Nonetheless, corals outplanted after only 1 month of rearing also had survivorship higher or comparable to other outplant methods currently being trialled, e.g. 25 % after 3 years in this study vs 8 % after 2.5 years for corals outplanted at 7 months old in Guest et al. (2014), 6 % after 2.7 years for corals outplanted at 5 months old in Humanes et al. (2021), and 32 % after 2.6 years for corals outplanted at 0.5 months in Chamberland et al. (2015). At 4 years post-deployment, outplant yield remained between 16 % and 36 %, considerably longer than the duration of the longest previous studies with sexually propagated Acropora (up to 2.7 years), which reported survival between 6 % and 47 % (Figure 7). These results suggest that long-term performance of CAPs may be higher than that of other devices if those were also surveyed after 4 years. Surviving colonies were found to be reproductively mature 4 years after outplant, demonstrating the feasibility of creating viable restored coral populations within half a decade. Realistic rehabilitation costs were comparable to or lower than other sexual propagation methods (i.e., clay tripods, direct seeding and coral plug-ins)(Chamberland et al., 2015; de la Cruz & Harrison, 2017; Guest et al., 2014) and could be further reduced with larger production scales (Humanes et al., 2021).

**Figure 7.**
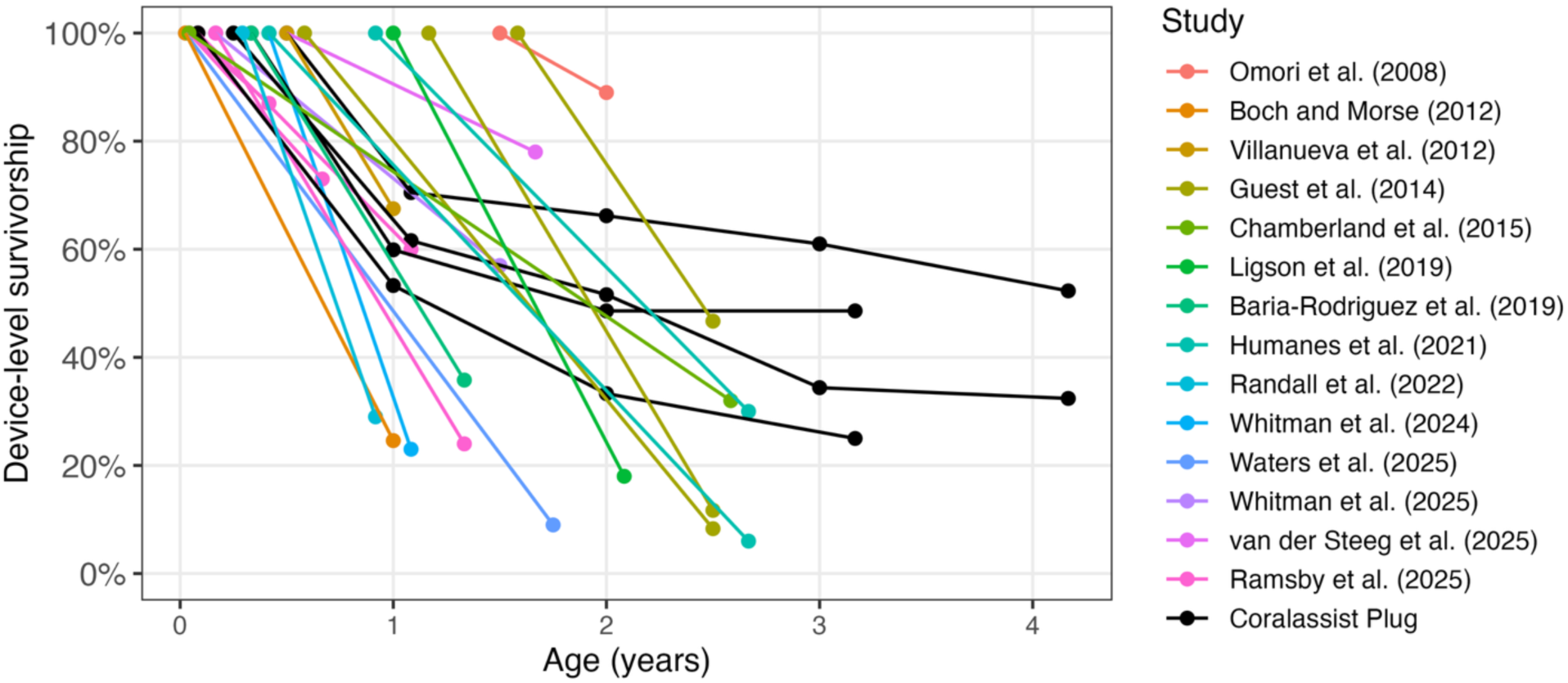
An overview of device-level survivorship of multiple studies that outplanted devices containing sexually reared *Acropora* to the reef. When multiple treatments, e.g. grazing deterrence, were compared, the best-performing method was included.

Fertilisation, larval development, settlement, and juvenile growth are the most vulnerable stages in the coral life cycle. Success during pre-settlement stages can be enhanced under controlled conditions, helping reduce major bottlenecks in coral propagation (Guest et al., 2010; Pollock et al., 2017). However, post-settlement mortality often remains high, both under controlled conditions (Craggs et al., 2019) and after deployment onto reefs (Humanes et al., 2021). The methods used to deploy corals onto degraded areas, as well as species-specific life-history traits (e.g., growth rates), strongly influence the survivorship of outplants, ultimately affecting the outcomes of rehabilitation interventions. Some of the processes affecting early survivorship may be density dependent (Doropoulos et al., 2017; Edwards et al., 2015), making it essential to determine optimal settlement densities. Optimising larval settlement density will be beneficial to maximising yield (Cameron & Harrison, 2020; Humanes et al., 2021), yet it is a challenging step in sexual coral propagation as it relies on the presence of specific cues that induce settlement (Randall et al., 2021). In artificial settlement devices, these cues are typically established through biological conditioning, whereby substrates are maintained in seawater for at least 6 weeks in the presence of crustose coralline algae (CCA). This process promotes the development of biofilms and CCA on device surfaces, both of which play a key role in influencing coral settlement (Heyward & Negri, 1999; Webster et al., 2004). However, the development of a CCA biofilm is impossible to standardise and control. This variability was evident in our study, where we observed substantial differences in CCA coverage across devices, resulting in significantly different settlement rates, despite all devices undergoing identical conditioning periods and treatment. Based on these observations, we recommend avoiding mixing devices with differing levels of conditioning within the same settlement tanks, as this will lead to uneven settlement densities among devices. Addressing the challenges associated with conditioning remains a crucial research area. Furthermore, achieving optimal conditioning with the appropriate CCA species may be specific to both coral species and geographic location (Abdul Wahab et al., 2023), and is essential for attaining the desired settlement densities.

Settlement densities will directly influence the number of corals per device, as well as their growth and survivorship. Here we determined optimal spat numbers at outplant for a range of nursery periods. Our results show that devices with a longer nursery rearing duration needed fewer corals at outplant to have at least 1 coral alive at 1 year old. This provides a useful guide for the optimal number of corals per device, which could help practitioners with decision-making. CAPs had a higher initial number of settled corals than the control devices in 2020, presumably because they had a larger surface area, and larvae may have preferred to settle within the provided microrefugia, as has been found in several other studies. However, in 2021 the number of spat was similar on both device types. Settlement at densities that are too low reduces the likelihood of long-term success and may render outplanting inefficient. Conversely, high densities can waste larvae and increase the risk of density-dependent mortality. High settlement densities have also been shown to promote the formation of chimeras (fusion of 2 or more larvae). Although information on the fitness of chimeras compared to single-genotype colonies across life stages is limited, chimeras grow faster than solitary juveniles during the first 3 months and may have higher survivorship during the first 8 to 12 months (Mendoza-Quiroz et al., 2026; Puill-Stephan et al., 2012; Raymundo & Maypa, 2004). This suggests that chimerism may be important in enhancing survival during vulnerable early life stages. However, it remains unclear whether chimeras persist in the long term (e.g., whether some genotypes are lost over time) or if they provide long-term benefits for reef rehabilitation, such as improved growth, survival, fecundity, or (heat) stress tolerance.

Increasing post-outplant survivorship will be crucial for successful implementation of sexual coral propagation in large-scale coral rehabilitation initiatives. High mortality rates of up to 80 % are commonly observed within the first year (Boch & Morse, 2012; Randall et al., 2022; Whitman et al., 2024), although longer nursery rearing durations and grazing deterrents can alleviate this bottleneck (Guest et al., 2014; Omori et al., 2008; van der Steeg et al., 2025)(Figure 7, see Appendix S1: Table S17). Note that the graph in Figure 7 shows (device-level) survivorship, not yield, which would likely be lower but is less commonly reported. For example, if 100 devices are outplanted, 10 are lost, and 45 contain live coral, this results in 50 % survivorship but 45 % yield. In this study, microrefugia on CAPs significantly improved post-outplant survivorship, leading to an 11 % increase in yield 4 years post-outplant compared to devices without microrefugia. Longer nursery durations of 6 months achieved 36 % yield 4 years post-outplant. The 6-month nursery duration increased yield almost twofold compared to the 1-month outplant at 3-years-old (47 % vs 24 %). The overall pattern of survivorship was similar across all outplant ages, with most mortality (23 % to 39 %) occurring within the first 3 months post-outplant, time after which survivorship stabilised, (Figure 5 A). This study demonstrates it is feasible to outplant corals as early as 1 month old and still attain device-level survivorship comparable to other studies (Figure 7).

High retention of settlement devices is crucial to maximise yield and impact of rehabilitation efforts. The retention rate of all devices was high at 92 % in the first 2 years but declined to 81 % and 71% at 3 and 4 years post-outplant, respectively. Although mortality rates stabilised after 2 years, the continued reduction in yield was mainly caused by the lower retention in the 3^rd^ and 4^th^ years. The outplant method was quick and reliable; however, the flexibility of the plastic push mount allowed the devices to move. This movement likely prevented corals from attaching to the reef substratum, with less than 50 % of surviving corals being self-attached after 2 years. The attachment of live coral tissue to the reef substratum is an important measure of outplant success. Corals attached to the reef are stabilised and much less likely to be dislodged and lost during storms or other disturbances (Guest et al., 2011). Detachment of the outplants indicates that a better method of attaching the CAPs to the reef should be developed. A future CAP could be designed for compatibility with the Coralclip®, bolts, or epoxy, which have demonstrated to provide a secure attachment and could reduce outplant time (Mendoza Quiroz et al., 2025; Suggett et al., 2019).

Enhanced growth rates allow coral spat to reach escape sizes faster and be recruited into the adult population earlier, but growth rates are highly variable across studies. For instance, 1- and 2-year-old corals in this study were smaller than *A. granulosa*, *A. millepora* and *A. verweyi* in the Philippines and *A. digitifera* in Palau (Baria-Rodriguez et al., 2019; Guest et al., 2014; Ligson et al., 2019; van der Steeg et al., 2025), yet larger than previous outplants of *A. hyacinthus* and *A. digitifera* in Palau (Boch & Morse, 2012; Humanes et al., 2021), and similar to *A. digitifera* in Australia (Whitman et al., 2025). Mature colonies were observed after 4 years, consistent with reports on the timing of reproductive onset for various *Acropora* species (Baria et al., 2012; Iwao et al., 2010; Lachs et al., 2026; Ligson & Cabaitan, 2021). Coral growth rates and outplant success more broadly are likely highly context-specific in both space and time.

Rehabilitation can play a crucial role in increasing recovery rates after disturbances and supporting coral populations and ecosystem functions (Fox et al., 2019; Lange et al., 2024; Montoya-Maya et al., 2016). Estimating costs can be a useful exercise to explore the most efficient methods and identify bottlenecks. Outplanting and monitoring were the largest expenses in the study, but in the realistic rehabilitation costing this shifted to outplanting and device costs, highlighting the need for efficient outplant methods and lower CAP costs. The estimated cost per coral at 2 years old or more for the experiment was lower compared to some studies (Guest et al., 2014; Humanes et al., 2021), but higher than others (Baria-Rodriguez et al., 2019; Chamberland et al., 2015), whereas the realistic rehabilitation costs were lower than all experimental costs reported to date (Table 1, Appendix S1: Table S17, Appendix S2). However, the higher salaries and boat costs of Palau compared to, e.g., the Philippines should be considered. The 6-month outplant with extended nursery rearing resulted in the lowest cost per coral at the end of the study, driven by high survivorship, consistent with Guest et al. (2014). The cost per coral increased faster for the 3-month outplant than for the 6-month outplant (77 % vs 30 % between 2 and 3 years post-outplant), attributed to greater ongoing mortality rates in the 3-month outplant. The cost of outplanting corals to a large reef area with this method will still be considerable, although it can be reduced by upscaling these methods, increasing nursery stocking density, outplant efficiency, and outplant survivorship (Scott et al., 2024; Suggett et al., 2023). The value of ecological rehabilitation should not be measured solely by the price per outplanted coral, but rather by maintaining the ecological, cultural, and intrinsic values of coral reef ecosystems (McAfee et al., 2021; Suggett et al., 2023, 2024).

The CAPs successfully achieved high survivorship over multiple years for corals outplanted to the reef after short and long nursery rearing durations, demonstrating the benefits of microrefugia. Across ecological contexts, there are likely to be numerous coral reef conservation and rehabilitation options that are best suited. CAPs are best suited for wave-exposed systems with limited natural recruitment and high grazing pressure, where they can achieve considerably high yield. Direct attachment can play an important role in local rehabilitation interventions, especially where reefs are easily and regularly accessible, or where diver time is relatively inexpensive. Our results point to two strategies for optimising outcomes: 1) outplant earlier, but in larger quantities to offset early losses, or 2) rear corals longer before outplanting, e.g. to 6 months old or ∼ 2 cm diameter, which requires rearing facilities and a dedicated husbandry team but delivers higher yields. A medium rearing duration still requires investment in rearing capacity without matching the yield gains of long rearing. Where continuous outplanting is more logistically feasible (e.g., weekly outplants with a dive club), this can still produce positive results over time. CAPs can be deployed at a multiple-hectare scale for reef rehabilitation efforts to accelerate recovery and increase ecosystem functioning. Further, their integration with sexual propagation and selective breeding makes this approach well-suited to incorporating ecosystem resilience through increasing genetic variation and adaptive capacity to future climates into management strategies.

## Acknowledgements

We want to thank the Palau International Coral reef Center and all their staff for their unwavering support, including Arius Merep for his guidance in building the experimental set up. Boat operators Nelson Masang and Geory Mereb and volunteers and research assistants Jayven Tachibelmel, Ernesto Andres JR, LeahMarie Bukurou, Kailey Gabrian-Voorhees, Craig Johnson, Janna Leigh Randle, Jada Avesani, Daniel Cassidy, Pierce Brooks, Louw Claassens, Minye Kim, Sharon Koen, Neil Nixon, Matt Boyle, McGee Mereb, Harrison Umang, Trinity Misech, Alik Ulechong, Elsei Tellei, Christina Muller-Karanassos, Alex Ferrier-Loh, Ayako Hosaka, Melanie Oborski and Arne Veimo who helped with coral spawning monitoring, larval rearing, outplanting and monitoring. We also thank the Bureau of Marine Resources for donating the rabbitfish used in this study. The work presented here was funded by the European Research Council Horizon 2020 project CORALASSIST (Project number 725848) and the European Research Council Proof of Concept grant (Project number EP/Y015290/1) awarded to JRG. The presented work was conducted under the Palau National Marine Research permits RE-20-04, RE-21-02, RE-22-11, RE-23-04 and RE-24-XX.

## Open Research Statement

The data will be made available upon request.

