## Appendix S1 for "The CoralAssist Plug: a novel device that enhances outplant rate and survivorship of sexually propagated corals"

**Section S1.** Coral larval rearing detailed methodology
On April 6th, 2 days before the full moon, 3 colonies spawned synchronously in the same holding tank at 20:30h. The gametes were collected in a 100L tank with 0.2 µm UV-treated FSW and left to fertilise for 45 minutes whilst the number of released eggs was estimated by counting 6 replicate 20ml aliquots. The embryos were washed to remove excess sperm and eight 45L rearing tanks with 0.2 µm UV-treated FSW were stocked with 15,000 embryos at a density of not more than 1 per 3 ml. After 90 min, fertilisation rates were estimated by counting the number of fertilised eggs in 5 replicate 20 ml aliquots taken from the surface of the fertilisation tank. Approximately 50% water changes were performed twice daily starting 20 hours after fertilisation. When the larvae became motile after approximately 36 hours, gentle aeration and water movement were introduced using rigid airlines. Four days after spawning, larval survivorship was estimated for each tank by counting 8 replicate 50 ml aliquots, and larval densities were reduced in preparation for settlement so that there were an estimated 50 larvae per device in each settlement tank.

Nursery rearing tanks had a constant 50 µm FSW flow-through at ~4 L/min and 2 pumps (Hydor Koralia Nano 425 Circulation Pump/Powerhead) to ensure sufficient water circulation. Each tank was provided with 2 aquarium lights (Reef Brite, 48” 50/50 me and blue XHO LED) at a 12:12 h diurnal cycle reaching up to 200 µmol photons $m^{-2}$ $s^{-1}$. Fragments of the parental colonies were added to the tanks to encourage *Symbiodiniaceae* uptake. Micro herbivores were added to the tanks to control algal growth and reduce manual cleaning. Each tank was stocked with 10 captive-bred juvenile rabbitfish (*Siganus lineatus,* ~5 cm long, donated by the Bureau of Marine Resources in Palau), 100 cerithid snails (*Cerithium sp.,* ~5 mm) and numerous trochus snails (*Trochus niloticus,* ~2 mm, spawned in August 2019). The fish were fed pellets daily (Ocean Nutrition Formula Two, small pellets), whereas the corals were fed 3 times a week (Coral Fusion plankton powder, Aqua Core) following the manufacturer’s instructions. The water flow and pumps were turned off for 30 minutes at the start of feeding, and the plankton powder was hydrated in 500ml FSW before addition. The nursery tanks were siphoned every other week to remove detritus and silt from the bottom, and algae were manually removed when needed. To control anemone (*Aiptasia* sp.*)* growth, 1 filefish (*Acreichthys tomentosus*, ~5 cm) collected from the PICRC dock was added to each tank 4 months after spawning in 2020.

**Table S1**. Model data for the relationship of device and conditioning type on the number of corals per device over time. The effect of days after settlement (continuous), device type (2 levels), conditioning type (2 levels) as fixed effects and tank (8 levels) as random effect on the number of corals per device was estimated using the count data of the 2020 subset using a generalised linear mixed effects model (GLMM) with a negative binomial distribution. Significance codes: *** $p < 0.001$, ** $p < 0.01$, * $p < 0.05$, n.s. = not significant

Model formula: Corals ~ Days + Treatment * Conditioning + (1 | Tank)
Model distribution: Negative Binomial (nbinom2) with log link
Model fit statistics: AIC = 809.5, BIC = 829.4, log-Lik = –397.7, deviance = 795.5, residual df: 121.

| Random effects | Name | Variance | Std.Dev |
|---|---|---|---|
| Tank | (Intercept) | 0. 4799 | 0.6927 |

Number of observations: 128, groups: Tank, 8
Dispersion parameter for nbinom2 family: 1.23

Fixed effects estimates:

| Predictor | Estimate | Std. Error | z value | p-value | Significance |
|---|---|---|---|---|---|
| Intercept | 3.0871 | 0.3312 | 9.320 | <0.001 | *** |
| Days | -0.0105 | 0.0032 | -3.302 | 0.0010 | *** |
| Treatment: CoralAssist Plug (vs Control) | 0.7018 | 0.2557 | 2.745 | 0.0060 | ** |
| Conditioning: Biofilm (vs CCA) | -1.9561 | 0.2844 | -6.879 | <0.001 | *** |
| Treatment × Conditioning interaction | -0.4274 | 0.3861 | -1.107 | 0.2683 | n.s. |

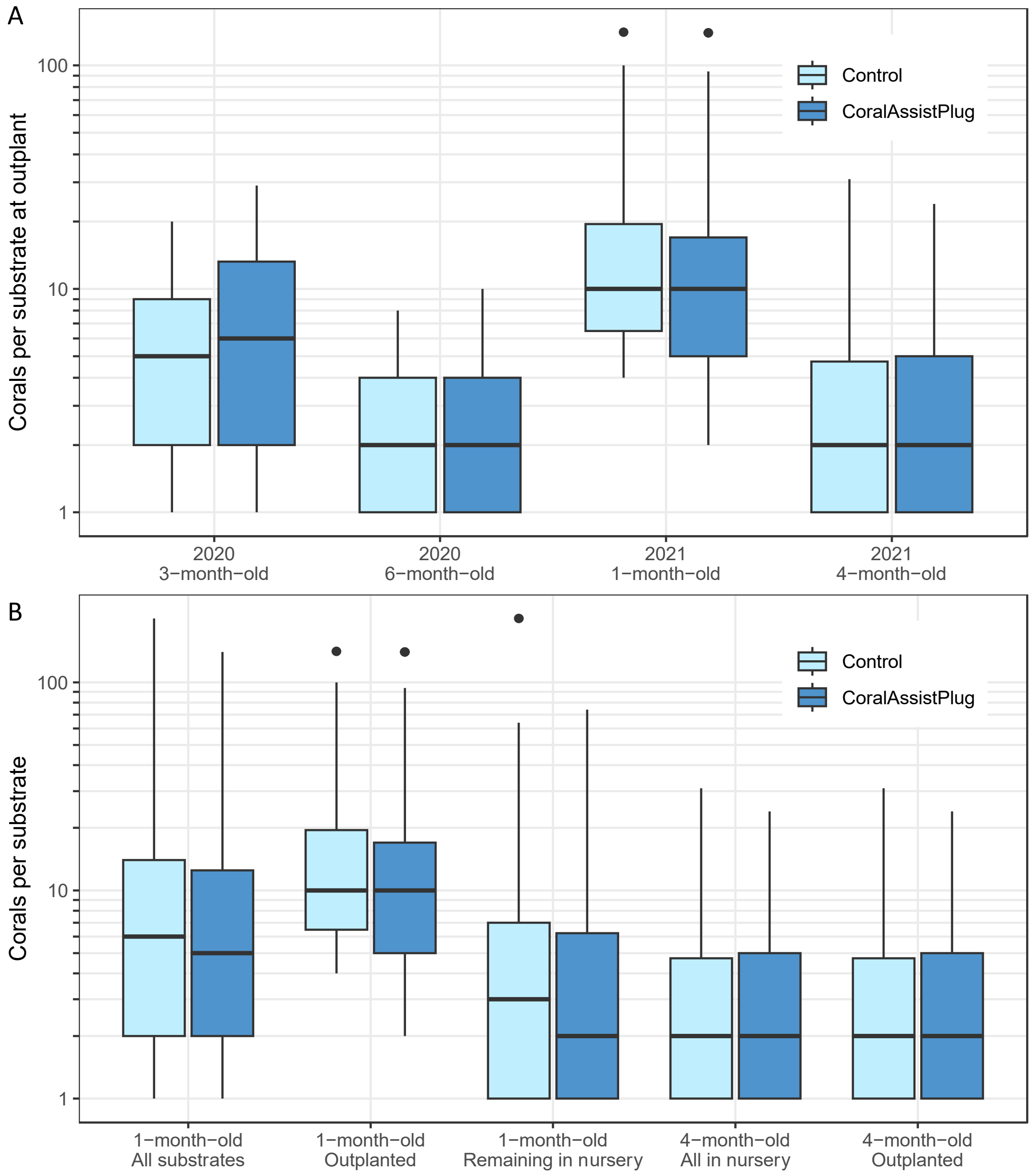


**Figure S1**. A) The number of corals per device at outplant on a logarithmic scale for all 4 outplant ages. B) The number of corals per device on a logarithmic scale in 2021, highlighting the priority of devices with at least 4 corals for the 1-month-old outplant, and the devices with a lower number of corals remaining in the nursery. The boxplots represent the interquartile range, the horizontal line in the box is the median and the whiskers show the minimum and maximum data values (1.5 * interquartile range) and the variability compared to the interquartile. The black dots are outliers outside of the whisker lines (> 1.5 * interquartile range from the median).

**Table S2**. Pairwise comparisons of the number of corals (log-transformed) per device at outplant between different outplant ages using a nonparametric Kruskal-Wallis test and a Bonferroni corrected pairwise Wilcoxon rank-sum test.

| | 2020 3-month | 2020 6-month | 2021 1-month |
|---|---|---|---|
| 2020 6-month | 1.7e-14 | - | - |
| 2021 1-month | 1.6e-10 | < 2e-16 | - |
| 2021 4-month | 2.7e-11 | 1.000 | < 2e-16 |

**Table S3**. Model parameters of GLMM for the effect of number of corals between outplant cohorts on the number of corals present after 1 year for only the CoralAssist Plugs. The effect of the log-transformed number of corals per device at outplant (continuous, fixed effect) and outplant cohort of only CAPs (4 levels, fixed effect) on the number of corals per device at 1-year-old was tested using a GLMM. We used a Poisson distribution, tested for overdispersion and ability to predict zeros, and fitted a random intercept for each transect (14 levels, random effect).

Model formula: No_Corals1Y ~ No_Corals_log + Cohort + (1 | Transect)
Generalised linear mixed model fit by maximum likelihood (Laplace Approximation) ['glmerMod'], Family: poisson ( log )
Model fit statistics: AIC = 798.8, BIC = 821.8, logLik = -393.4, deviance = 786.8, df.resid = 335.
Scaled residuals: Min = -1.7369, 1Q = -0.7648, Median = -0.0918, 3Q = 0.5066, Max = 3.6711.

Random effects:

| Groups | Name | Variance | Std.Dev |
|---|---|---|---|
| Transect | (Intercept) | 0 | 0 |

Number of obs: 341, groups: Transect, 14

Fixed effects:

| | Estimate | Std. Error | z value | Pr(>\|z\|) |
|---|---|---|---|---|
| (Intercept) | -1.07351 | 0.17132 | -6.266 | 3.70e-10 *** |
| No_Corals_log | 0.52078 | 0.06277 | 8.296 | < 2e-16 *** |
| Cohort2020\n6-month-old | 0.63239 | 0.15768 | 4.011 | 6.05e-05 *** |
| Cohort2021\n1-month-old | -0.39585 | 0.16690 | -2.372 | 0.0177 * |
| Cohort2021\n4-month-old | 0.25042 | 0.18581 | 1.348 | 0.1777 |

Signif. codes: *** p < 0.001, ** p < 0.01, * p < 0.05

Correlation of Fixed Effects:

| | (Intr) | N_Crl_ | C_2020_6 | C_2021_1 |
|---|---|---|---|---|
| No_Corls_lg | -0.796 | | | |
| Chrt_2020_6 | -0.743 | 0.434 | | |
| Chrt_2021_1 | -0.178 | -0.248 | 0.301 | |
| Chrt_2021_4 | -0.586 | 0.312 | 0.502 | 0.269 |

| contrast | estimate | SE | df | z.ratio | p.value |
|---|---|---|---|---|---|
| (2020\n3-month-old) - (2020\n6-month-old) | -0.632 | 0.158 | Inf | -4.011 | 0.0004 *** |
| (2020\n3-month-old) - (2021\n1-month-old) | 0.396 | 0.167 | Inf | 2.372 | 0.0826 |
| (2020\n3-month-old) - (2021\n4-month-old) | -0.250 | 0.186 | Inf | -1.348 | 0.5324 |
| (2020\n6-month-old) - (2021\n1-month-old) | 1.028 | 0.192 | Inf | 5.354 | <.0001 *** |
| (2020\n6-month-old) - (2021\n4-month-old) | 0.382 | 0.173 | Inf | 2.207 | 0.1212 |
| (2021\n1-month-old) - (2021\n4-month-old) | -0.646 | 0.214 | Inf | -3.024 | 0.0133 * |

Results are given on the log (not the response) scale.
P value adjustment: tukey method for comparing a family of 4 estimates.

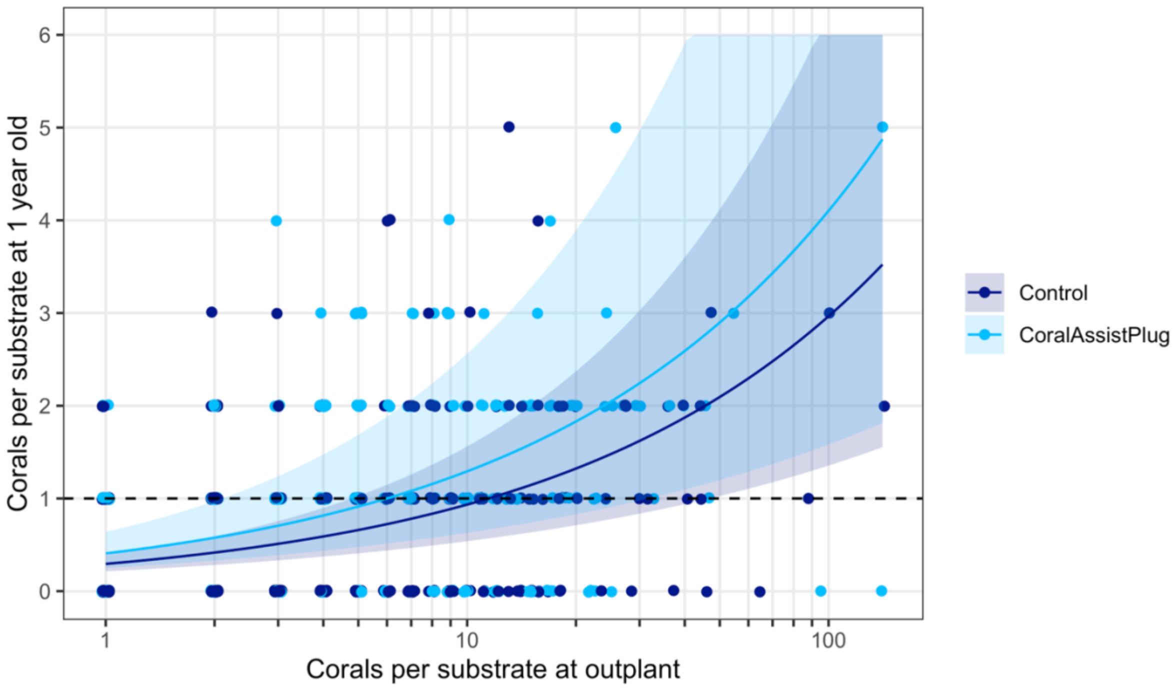


**Figure S2**. The relationship between corals per device at outplant and corals per device at 1year old, grouped device type, derived from a GLMM with Poisson distribution. The original data is represented with a jitter to prevent overlap, and the shaded areas represent 95% confidence intervals. The dashed line represents 1 coral alive at 1 year old.

**Table S4**. Model parameters of the GLMM for the effect of number of corals at outplant on the number of corals present after 1 year for the CoralAssist Plugs and control devices. The effect of the log-transformed number of corals per device at outplant (continuous, fixed effect) and device type (2 levels, fixed effect) on the number of corals per device at 1-year-old was tested using a GLMM. We used a Poisson distribution, tested for overdispersion and ability to predict zeros, and fitted a random intercept for each transect (14 levels, random effect).

Generalised linear mixed model fit by maximum likelihood (Laplace Approximation) ['glmerMod'], Family: poisson ( log )
Formula: No_Corals1Y ~ No_Corals_log + Treatment + (1 | Transect)
Model fit statistics: AIC = 1549.0, BIC = 1567.2, logLik = -770.5, deviance = 1541.0, df.resid = 694.
Scaled residuals: Min = -1.8863, 1Q = -0.7534, Median = -0.2293, 3Q = 0.6101, Max = 3.8979.

| Random effects | Name | Variance | Std.Dev |
|---|---|---|---|
| Transect | (Intercept) | 0.1341 | 0.3661 |

Number of obs: 698, groups: Transect, 14

| Fixed effects | Estimate | Std. Error | z value | Pr(>\|z\|) |
|---|---|---|---|---|
| (Intercept) | -1.22315 | 0.14842 | -8.241 | < 2e-16 *** |
| No_Corals_log | 0.50162 | 0.05144 | 9.752 | < 2e-16 *** |
| TreatmentCoralAssistPlug | 0.32472 | 0.08620 | 3.767 | 0.000165 *** |

Signif. codes: *** p < 0.001, ** p < 0.01, * p < 0.05

| Correlation of Fixed Effects | (Intercept) | No_Corals_log |
|---|---|---|
| No_Corals_log | -0.609 | |
| TreatmentCoralAssistPlug | -0.321 | -0.017 |

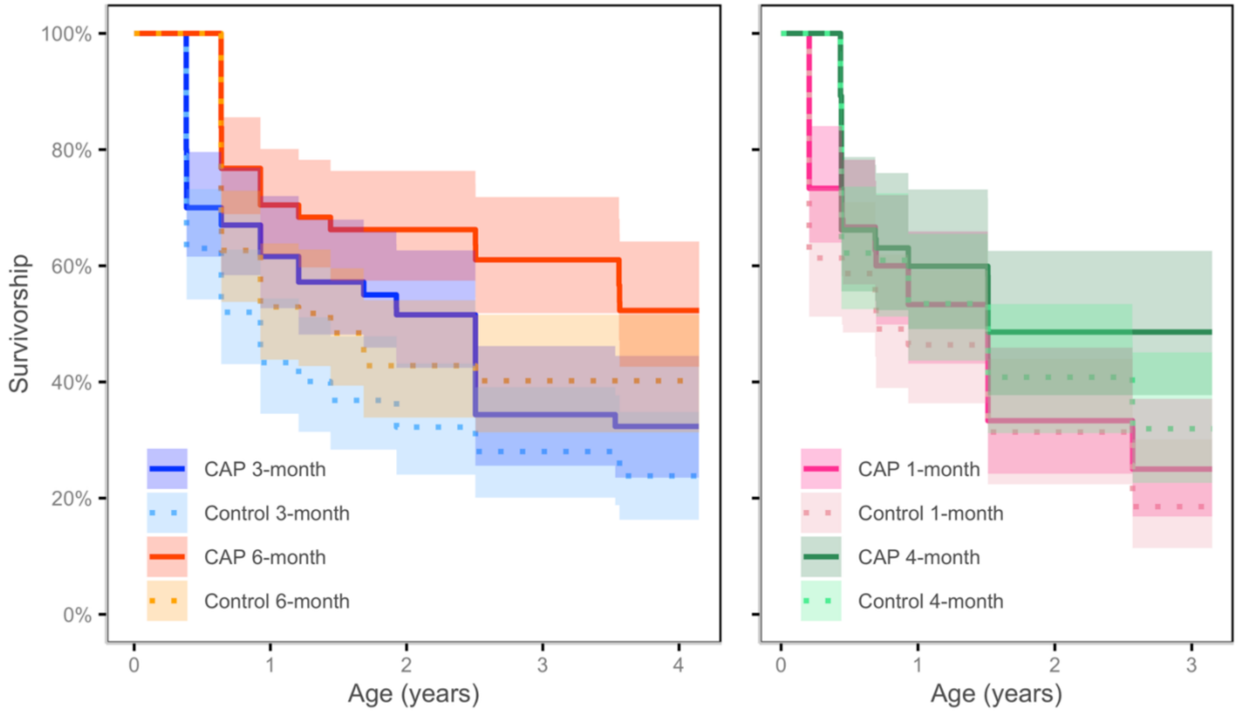


**Figure S3**. Survival curves with 95 % confidence intervals of CoralAssist Plug (CAP) and control devices outplanted to the reef after varying nursery periods in 2020 and 2021. Day=0 is the day of spawning. Solid lines are the CAP, dashed lines are the smooth control devices. Devices outplanted after a 3-month nursery period are depicted in blue, 6-months in orange and 1-month in pink. In 2020 n=100 for each device type, in 2021 n = 75 for both 1-month outplants and for the 3-month outplant n=66 for the CAPs and n = 82 for the control.

**Table S5**. Mean survival ages of CoralAssist Plug (CAP) and control devices outplanted after different nursery durations.

| Year | Treatment | Cohort | N | Events | R mean | SE (R mean) | median | 0.95 LCL | 0.95 UCL |
|---|---|---|---|---|---|---|---|---|---|
| 2020 | CAP | 3-month | 100 | 61 | 798* | 59.4 | 915 | 441 | 915 |
| 2020 | Control | 3-month | 100 | 72 | 625* | 58.3 | 338 | 234 | 526 |
| 2020 | CAP | 6-month | 100 | 43 | 1049* | 57.8 | NA | 1300 | NA |
| 2020 | Control | 6-month | 100 | 57 | 810* | 61.0 | 526 | 338 | NA |
| 2021 | CAP | 1-month | 75 | 56 | 546** | 49.2 | 552 | 254 | 552 |
| 2021 | Control | 1-month | 75 | 60 | 483** | 49.6 | 254 | 162 | 552 |
| 2021 | CAP | 4-month | 66 | 34 | 694** | 57.1 | 552 | 340 | NA |
| 2021 | Control | 4-month | 82 | 52 | 610** | 48.5 | 552 | 340 | 937 |

* restricted mean with upper limit = 1515

** restricted mean with upper limit = 1150

**Table S6**. Pairwise comparisons of survival curves using the Log-Rank test with Bonferroni-adjusted p-values. Abbreviations: CAP = CoralAssist Plug; Ctrl = Control; 1m/3m/4m/6m = 1-/3-/4-/6-month-old cohorts. Significance codes: *** $p < 0.001$, ** $p < 0.01$, * $p < 0.05$

| | 2020 CAP 3m | 2020 Ctrl 3m | 2020 CAP 6m | 2020 Ctrl 6m | 2021 CAP 1m | 2021 Ctrl 1m | 2021 CAP 4m |
|---|---|---|---|---|---|---|---|
| 2020 Ctrl 3m | 1.0000 | – | | | | | |
| 2020 CAP 6m | 0.0435 * | 1.7e-05 *** | – | | | | |
| 2020 Ctrl 6m | 1.0000 | 0.1314 | 0.4919 | – | | | |
| 2021 CAP 1m | 1.0000 | 1.0000 | 7.7e-05 *** | 0.9902 | – | | |
| 2021 Ctrl 1m | 0.0988 | 1.0000 | 2.8e-07 *** | 0.0328 * | 1.0000 | – | |
| 2021 CAP 4m | 1.0000 | 0.1572 | 0.7236 | 1.0000 | 0.4177 | 0.0179 * | – |
| 2021 Ctrl 4m | 1.0000 | 1.0000 | 0.0018 ** | 1.0000 | 1.0000 | 0.8610 | 1.0000 |

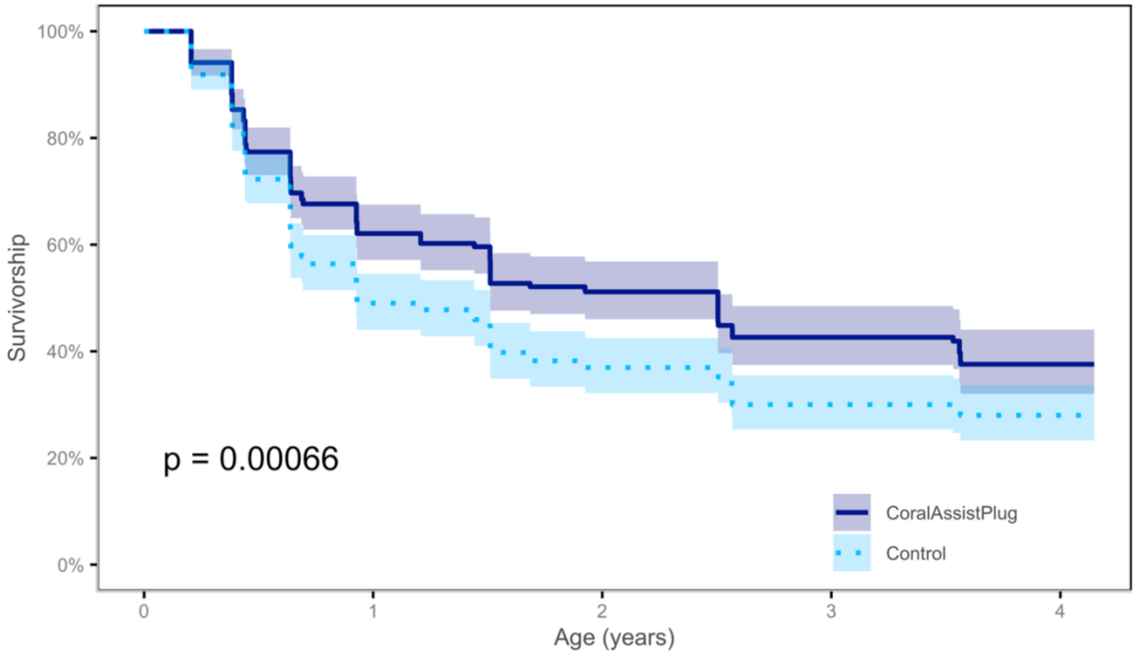


**Figure S4**. Survival curves with 95 % confidence intervals of all 4 outplants combined and grouped by device type; CoralAssist Plug (CAP) and control. Day=0 is the day of spawning, n=341 for CAPs and n=357 for the control.

**Table S7**. Mean survival ages of CoralAssist Plug (CAP) and control devices, restricted mean survival with upper limit = 1515 days.

| Treatment | N | Events | R mean | SE (R mean) | median | 0.95 LCL | 0.95 UCL |
|---|---|---|---|---|---|---|---|
| CAP | 341 | 193 | 845 | 33.0 | 915 | 552 | 938 |
| Control | 357 | 241 | 681 | 31.4 | 340 | 338 | 552 |

**Table S8.** Model data for the relationship of device type, outplant cohort and transect on the retention of devices outplanted in 2020. The effect of device type (2 levels, fixed effect), outplant cohort (2 levels, fixed effect) and transect (8 levels, random effect) on the retention was estimated using a generalised linear mixed effects model (GLMM) with a binominal distribution with a logit-link function using glmer in the lme4 package in R. Significance codes: *** p < 0.001, ** p < 0.01, * p < 0.05.

Model formula: Retention ~ Treatment + Outplant + (1 | Transect), Data: Retention2020
Model distribution: binominal with logit link
Model fit statistics: AIC = 491.2, BIC = 507.2, logLik = -241.6, deviance = 483.2, residual df = 396.
Scaled residuals: Min = -1.7624, 1Q = -1.3522, Median = 0.6027, 3Q = 0.6962, Max = 0.7395.

| Random effects | Name | Variance | Std.Dev |
|---|---|---|---|
| Transect | (Intercept) | 0 | 0 |

Number of observations: 400, groups: Transect; 8

| Fixed effect | Estimate | Std. Error | z value | p-value | Significance |
|---|---|---|---|---|---|
| Intercept (Control, 3-month) | 1.0127 | 0.1942 | 5.214 | 1.84e-07 | *** |
| Treatment: CoralAssist Plug (vs Control) | -0.4091 | 0.2205 | -1.855 | 0.0636 | |
| Outplant: 6 month (vs 3-month) | 0.1207 | 0.2198 | 0.549 | 0.5829 | |

| Correlation of fixed effects | (Intercept) | Treatment: CoralAssist Plug |
|---|---|---|
| Treatment: CorAlassist Plug | -0.612 | |
| Outplant: 6 month | -0.549 | -0.005 |

**Table S9.** Model data for the relationship of device type, outplant cohort and transect on the retention of devices outplanted in 2021. The effect of device type (2 levels, fixed effect), outplant cohort (2 levels, fixed effect) and transect (6 levels, random effect) on the retention was estimated using a generalised linear mixed effects model (GLMM) with a binominal distribution with a logit-link function using glmer in the lme4 package in R. Significance codes: *** p < 0.001, ** p < 0.01, * p < 0.05.

Model formula: Retention ~ Treatment + Outplant + (1 | Transect), Data: Retention2021
Model distribution: binominal with logit link
Model fit statistics: AIC = 222.6, BIC = 237.4, logLik = -107.3, deviance = 214.6, residual df = 294.
Scaled residuals: Min = -3.5414, 1Q = 0.2824, Median = 0.3070, 3Q = 0.4233, Max = 0.4603.

| Random effects | Name | Variance | Std.Dev |
|---|---|---|---|
| Transect | (Intercept) | 0 | 0 |

Number of observations: 298, groups: Transect; 6

| Fixed effect | Estimate | Std. Error | z value | p-value | Significance |
|---|---|---|---|---|---|
| Intercept (Control, 1-month) | 2.5290 | 0.3572 | 7.081 | 1.43e-12 | *** |
| Treatment: Coralassist Plug (vs Control) | -0.1675 | 0.3591 | -0.466 | 0.641 | |
| Outplant: 4 month (vs 1-month) | -0.8098 | 0.3753 | -2.157 | 0.031 | * |

| Correlation of fixed effects | (Intercept) | Treatment: CoralAssist Plug |
|---|---|---|
| Treatment: CoralAssist Plug | -0.538 | |
| Outplant: 4 month | -0.708 | -0.059 |

**Table S10.** Model data for the relationship of device type, outplant cohort and transect on device yield (retained and containing ≥ 1 live coral) of devices outplanted in 2020. The effect of device type (2 levels, fixed effect), outplant cohort (2 levels, fixed effect) and transect (8 levels, random effect) on device yield was estimated using a generalised linear mixed effects model (GLMM) with a binominal distribution with a logit-link function using glmer in the lme4 package in R. Significance codes: *** p < 0.001, ** p < 0.01, * p < 0.05.

Model formula: Yield ~ Treatment + Outplant + (1 | Transect), Data: Yield2020
Model distribution: binominal with logit link
Model fit statistics: AIC = 430.4, BIC = 446.4, logLik = -211.2, deviance = 422.4, residual df = 396.
Scaled residuals: Min = -0.7793, 1Q = -0.5929, Median = -0.4706, 3Q = -0.3872, Max = 2.5823.

| Random effects | Name | Variance | Std.Dev |
|---|---|---|---|
| Transect | (Intercept) | 0.07333 | 0.2708 |

Number of observations: 400, groups: Transect; 8

| Fixed effect | Estimate | Std. Error | z value | p-value | Significance |
|---|---|---|---|---|---|
| Intercept (Control, 3-month) | -1.7920 | 0.2700 | -6.636 | 3.22e-11 | *** |
| Treatment: CoralAssist Plug (vs Control) | 0.2909 | 0.2417 | 1.203 | 0.2288 | |
| Outplant: 6 month (vs 3-month) | 0.8008 | 0.3124 | 2.563 | 0.0104 | * |

| Correlation of fixed effects | (Intercept) | Treatment: CoralAssist Plug |
|---|---|---|
| Treatment: CoralAssist Plug | -0.492 | |
| Outplant: 6 month | -0.659 | -0.016 |

**Table S11.** Model data for the relationship of device type, outplant cohort and transect on device yield (retained and containing ≥ 1 live coral) of devices outplanted in 2021. The effect of device type (2 levels, fixed effect), outplant cohort (2 levels, fixed effect) and transect (6 levels, random effect) on device yield was estimated using a generalised linear mixed effects model (GLMM) with a binominal distribution with a logit-link function using glmer in the lme4 package in R. Significance codes: *** p < 0.001, ** p < 0.01, * p < 0.05.

Model formula: Yield ~ Treatment + Outplant + (1 | Transect), Data: Yield2021
Model distribution: binominal with logit link
Model fit statistics: AIC = 333.0, BIC = 347.8, logLik = -162.5, deviance = 325.0, residual df = 294.
Scaled residuals: Min = -0.7317, 1Q = -0.5822, Median = -0.5489, 3Q = -0.4367, Max = 2.2897.

| Random effects | Name | Variance | Std.Dev |
|---|---|---|---|
| Transect | (Intercept) | 0 | 0 |

Number of observations: 298, groups: Transect; 6

| Fixed effect | Estimate | Std. Error | z value | p-value | Significance |
|---|---|---|---|---|---|
| Intercept (Control, 1-month) | -1.6568 | 0.2586 | -6.406 | 1.49e-10 | *** |
| Treatment: CoralAssist Plug (vs Control) | 0.5749 | 0.2743 | 2.096 | 0.0361 | * |
| Outplant: 4 month (vs 1-month) | 0.4572 | 0.2747 | 1.664 | 0.0961 | |

| Correlation of fixed effects | (Intercept) | Treatment: CoralAssist Plug |
|---|---|---|
| Treatment: CoralAssist Plug | -0.620 | |
| Outplant: 4 month | -0.628 | -0.077 |

**Table S12.** Model data for the relationship of device type, outplant cohort and transect on the self-attachment of corals outplanted in 2020. The effect of device type (2 levels, fixed effect), outplant cohort (2 levels, fixed effect) and transect (8 levels, random effect) on self-attachment was estimated using a generalised linear mixed effects model (GLMM) with a binominal distribution with a logit-link function using glmer in the lme4 package in R. Significance codes: *** p < 0.001, ** p < 0.01, * p < 0.05.

Model formula: Self-attachment ~ Treatment + Outplant + (1 | Transect), Data: Attachment2020
Model distribution: binominal with logit link
Model fit statistics: AIC = 76.5, BIC = 86.7, logLik = -34.2, deviance = 68.5, residual df = 90.
Scaled residuals: Min = -4.5631, 1Q = 0.1195, Median = 0.3761, 3Q = 0.4071, Max = 0.7357.

| Random effects | Name | Variance | Std.Dev |
|---|---|---|---|
| Transect | (Intercept) | 0.04285 | 0.207 |

Number of observations: 94, groups: Transect; 8

| Fixed effect | Estimate | Std. Error | z value | p-value | Significance |
|---|---|---|---|---|---|
| Intercept (Control, 3-month) | 3.0673 | 1.0422 | 2.943 | 0.00325 | ** |
| Treatment: CoralAssist Plug (vs Control) | 1.1837 | 0.6379 | 1.856 | 0.06348 | |
| Outplant: 6 month (vs 3-month) | -2.3470 | 1.0893 | -2.155 | 0.03119 | * |

| Correlation of fixed effects | (Intercept) | Treatment: CoralAssist Plug |
|---|---|---|
| Treatment: CoralAssist Plug | -0.125 | |
| Outplant: 6 month | -0.016 | -0.119 |

**Table S13.** Model data for the relationship of device type, outplant cohort and transect on the self-attachment of corals outplanted in 2021. The effect of device type (2 levels, fixed effect), outplant cohort (2 levels, fixed effect) and transect (6 levels, random effect) on self-attachment was estimated using a generalised linear mixed effects model (GLMM) with a binominal distribution with a logit-link function using glmer in the lme4 package in R. Significance codes: *** p < 0.001, ** p < 0.01, * p < 0.05.

Model formula: Sefl-attachment ~ Treatment + Outplant + (1 | Transect), Data: Attachment2021
Model distribution: binominal with logit link
Model fit statistics: AIC = 83.2, BIC = 92.4, logLik = -37.6, deviance = 75.2, residual df = 69.
Scaled residuals: Min = -2.4910, 1Q = 0.4014, Median = 0.4882, 3Q = 0.5454, Max = 0.6632.

| Random effects | Name | Variance | Std.Dev |
|---|---|---|---|
| Transect | (Intercept) | 0 | 0 |

Number of observations: 73, groups: Transect; 6

| Fixed effect | Estimate | Std. Error | z value | p-value | Significance |
|---|---|---|---|---|---|
| Intercept (Control, 1-month) | 1.4342 | 0.5717 | 2.509 | 0.0121 | * |
| Treatment: Coralassist Plug (vs Control) | -0.6128 | 0.6032 | -1.016 | 0.3097 | |
| Outplant: 4 month (vs 1-month) | 0.3912 | 0.5725 | 0.683 | 0.4944 | |

| Correlation of fixed effects | (Intercept) | Treatment: CoralAssist Plug |
|---|---|---|
| Treatment: CoralAssist Plug | -0.690 | |
| Outplant: 4 month | -0.516 | -0.010 |

**Table S14**. Model data for the relationship of device type and outplant cohort on the geometric mean diameter (GMD) of the largest coral on a device at 4-years-old from corals from 2020. The effect of device type (2 levels, fixed effect), outplant cohort (2 levels, fixed effect) and transect (8 levels, random effect) on the GMD was estimated using a generalised linear mixed effects model (GLMM) with a lognormal error distribution using the glmmTMB package in R. Significance codes: *** p < 0.001, ** p < 0.01, * p < 0.05.

Model formula: GMD_largest ~ Treatment + OutplantCohort + (1 | Transect)
Model distribution: gaussian with log link
Model fit statistics: AIC = 461.3, BIC = 474.0, logLik = -225.6, deviance = 451.3, residual df = 89.

| Random effects | Name | Variance | Std.Dev |
|---|---|---|---|
| Transect | (Intercept) | 0.01396 | 0.1182 |
| Residual | | 6.43655 | 2.5370 |

Number of obs: 94, groups: Transect, 8
Dispersion estimate for gaussian family (sigma^2): 6.44

| Fixed effect | Estimate | Std. Error | z value | p-value | Significance |
|---|---|---|---|---|---|
| Intercept (Control, 3-month) | 2.3188 | 0.0787 | 29.449 | <0.001 | *** |
| Treatment: CoralAssist Plug (vs Control) | 0.0120 | 0.0555 | 0.216 | 0.829 | |
| Outplant cohort: 6-month (vs 3-month) | -0.1075 | 0.1020 | -1.055 | 0.292 | |

Planned pairwise contrasts (ratios of means, log scale tested)

| Contrast | Ratio | SE | df | t-ratio | p-value |
|---|---|---|---|---|---|
| Control 3-m / CoralAssist Plug 3-m | 0.988 | 0.0549 | 89 | -0.216 | 0.996 |
| Control 3-m / Control 6-m | 1.114 | 0.1140 | 89 | 1.055 | 0.718 |
| Control 3-m / CoralAssist Plug 6-m | 1.100 | 0.1240 | 89 | 0.848 | 0.831 |
| CoralAssist Plug 3-m / Control 6-m | 1.127 | 0.1350 | 89 | 1.000 | 0.750 |
| CoralAssist Plug 3-m / CoralAssist Plug 6-m | 1.114 | 0.1140 | 89 | 1.055 | 0.718 |
| Control 6-m / CoralAssist Plug 6-m | 0.988 | 0.0549 | 89 | -0.216 | 0.996 |

Means and confidence intervals are back-transformed from the log scale. Pairwise contrasts were adjusted with Tukey’s method.

**Table S15**. Model data for the relationship of device type and outplant cohort on the geometric mean diameter (GMD) of the largest coral on a device at 3-years-old from corals from 2021. The effect of device type (2 levels, fixed effect), outplant cohort (2 levels, fixed effect) and transect (6 levels, random effect) on the GMD was estimated using a generalised linear mixed effects model (GLMM) with a lognormal error distribution using the glmmTMB package in R. Significance codes: *** p < 0.001, ** p < 0.01, * p < 0.05.

Model formula: GMD_largest ~ Treatment + OutplantCohort + (1 | Transect)
Model distribution: gaussian with log link
Model fit statistics: AIC = 314.9, BIC = 326.4, logLik = -152.5, deviance = 304.9, residual df = 68.

| Random effects | Name | Variance | Std.Dev |
|---|---|---|---|
| Transect | (Intercept) | 0.01303 | 0.1142 |
| Residual | | 3.51752 | 1.8755 |

Number of obs: 73, groups: Transect, 6
Dispersion estimate for gaussian family (sigma^2): 3.52

| Fixed effect | Estimate | Std. Error | z value | p-value | Significance |
|---|---|---|---|---|---|
| Intercept (Control, 1-month) | 1.76480 | 0.09720 | 18.156 | <2e-16 | *** |
| Treatment: CoralAssist Plug (vs Control) | -0.06258 | 0.07167 | -0.873 | 0.383 | |
| Outplant cohort: 4-month (vs 1-month) | 0.16358 | 0.11931 | 1.371 | 0.170 | |

Planned pairwise contrasts (ratios of means, log scale tested)

| Contrast | Ratio | SE | df | t-ratio | p-value |
|---|---|---|---|---|---|
| Control 1-m / CoralAssist Plug 1-m | 0.354 | 0.408 | 68 | 0.868 | 0.8212 |
| Control 1-m / Control 4-m | -1.038 | 0.755 | 68 | -1.374 | 0.5199 |
| Control 1-m / CoralAssist Plug 4-m | -0.621 | 0.849 | 68 | -0.731 | 0.8842 |
| CoralAssist Plug 1-m / Control 4-m | -1.392 | 0.862 | 68 | -1.615 | 0.3769 |
| CoralAssist Plug 1-m / CoralAssist Plug 4-m | -0.975 | 0.709 | 68 | -1.376 | 0.5187 |
| Control 4-m / CoralAssist Plug 4-m | 0.417 | 0.481 | 68 | 0.868 | 0.8213 |

Means and confidence intervals are back-transformed from the log scale. Pairwise contrasts were adjusted with Tukey’s method.

**Table S16**. Model data for the relationship of device type and outplant cohort on the geometric mean diameter (GMD) of the largest coral on a device at 3-years-old from corals from 2020 and 2021. The effect of device type (2 levels, fixed effect), outplant cohort (4 levels, fixed effect) and transect (14 levels, random effect) on the GMD was estimated using a generalised linear mixed effects model (GLMM) with a lognormal error distribution using the glmmTMB package in R. Significance codes: *** p < 0.001, ** p < 0.01, * p < 0.05.

Model formula: GMD_largest ~ Treatment + OutplantCohort + (1 | Transect) + (1 | Year)
Model distribution: gaussian with log link
Model fit statistics: AIC = 809.5, BIC = 835.7, logLik = -396.7, deviance = 793.5, residual df = 189.

| Random effects | Name | Variance | Std.Dev |
|---|---|---|---|
| Transect | (Intercept) | 0.004879 | 0.06985 |
| Year | (Intercept) | <0.000001 | <0.000001 |
| Residual | | 3.174388 | 1.78168 |

Number of obs: 197, groups: Transect: 14, Year: 2
Dispersion estimate for gaussian family (sigma^2): 3.17

| Fixed effect | Estimate | Std. Error | z value | p-value | Significance |
|---|---|---|---|---|---|
| Intercept (Control, 1-month) | 1.748887 | 0.07440 | 23.506 | <2e-16 | *** |
| Treatment: CoralAssist Plug (vs Control) | -0.02803 | 0.04712 | -0.595 | 0.5520 | |
| Outplant cohort: 3-month (vs 1-month) | -0.15610 | 0.09506 | -1.642 | 0.1006 | |
| Outplant cohort: 4-month (vs 1-month) | 0.16106 | 0.09066 | 1.776 | 0.0757 | |
| Outplant cohort: 6-month (vs 1-month) | -0.11164 | 0.08820 | -1.266 | 0.2056 | |

Planned pairwise contrasts (ratios of means, log scale tested)

| Contrast | Ratio | SE | df | t-ratio | p-value | Significance |
|---|---|---|---|---|---|---|
| Control 1-m / CoralAssist Plug 1-m | 0.1589 | 0.268 | 190 | 0.593 | 0.9989 | |
| Control 1-m / Control 3-m | 0.8307 | 0.512 | 190 | 1.623 | 0.7359 | |
| Control 1-m / CoralAssist Plug 3-m | 0.9666 | 0.565 | 190 | 1.711 | 0.6803 | |
| Control 1-m / Control 4-m | -1.0045 | 0.561 | 190 | -1.790 | 0.6276 | |
| Control 1-m / CoralAssist Plug 4-m | -0.8179 | 0.621 | 190 | -1.318 | 0.8911 | |
| Control 1-m / Control 6-m | 0.6072 | 0.487 | 190 | 1.246 | 0.9170 | |
| Control 1-m / CoralAssist Plug 6-m | 0.7493 | 0.542 | 190 | 1.384 | 0.8638 | |
| CoralAssist Plug 1-m / Control 3-m | 0.6719 | 0.559 | 190 | 1.202 | 0.9306 | |
| CoralAssist Plug 1-m / CoralAssist Plug 3-m | 0.8078 | 0.497 | 190 | 1.624 | 0.7354 | |
| CoralAssist Plug 1-m / Control 4-m | -1.1634 | 0.629 | 190 | -1.851 | 0.5862 | |
| CoralAssist Plug 1-m / CoralAssist Plug 4-m | -0.9768 | 0.545 | 190 | -1.792 | 0.6261 | |
| CoralAssist Plug 1-m / Control 6-m | 0.4483 | 0.546 | 190 | 0.822 | 0.9917 | |
| CoralAssist Plug 1-m / CoralAssist Plug 6-m | 0.5904 | 0.474 | 190 | 1.244 | 0.9174 | |
| Control 3-m / CoralAssist Plug 3-m | 0.1359 | 0.229 | 190 | 0.594 | 0.9989 | |
| Control 3-m / Control 4-m | -1.8353 | 0.508 | 190 | -3.613 | 0.0091 | ** |

| Contrast | Ratio | SE | df | t-ratio | p-value | Significance |
|---|---|---|---|---|---|---|
| Control 3-m / CoralAssist Plug 4-m | -1.6486 | 0.560 | 190 | -2.943 | 0.0700 | |
| Control 3-m / Control 6-m | -0.2235 | 0.422 | 190 | -0.530 | 0.9995 | |
| Control 3-m / CoralAssist Plug 6-m | -0.0815 | 0.472 | 190 | -0.173 | 1.0000 | |
| CoralAssist Plug 3-m / Control 4-m | -1.9712 | 0.575 | 190 | -3.431 | 0.0166 | * |
| CoralAssist Plug 3-m / CoralAssist Plug 4-m | -1.7845 | 0.492 | 190 | -3.627 | 0.0087 | ** |
| CoralAssist Plug 3-m / Control 6-m | -0.3595 | 0.483 | 190 | -0.745 | 0.9955 | |
| CoralAssist Plug 3-m / CoralAssist Plug 6-m | -0.2174 | 0.410 | 190 | -0.530 | 0.9995 | |
| Control 4-m / CoralAssist Plug 4-m | 0.1866 | 0.315 | 190 | 0.593 | 0.9989 | |
| Control 4-m / Control 6-m | 1.6117 | 0.482 | 190 | 3.342 | 0.0220 | * |
| Control 4-m / CoralAssist Plug 6-m | 1.7538 | 0.551 | 190 | 3.182 | 0.0357 | |
| CoralAssist Plug 4-m / Control 6-m | 1.4251 | 0.547 | 190 | 2.604 | 0.1609 | |
| CoralAssist Plug 4-m / CoralAssist Plug 6-m | 1.5672 | 0.469 | 190 | 3.342 | 0.0220 | * |
| Control 6-m / CoralAssist Plug 6-m | 0.1421 | 0.240 | 190 | 0.593 | 0.9989 | |

Means and confidence intervals are back-transformed from the log scale. Pairwise contrasts were adjusted with Tukey’s method.

**Table S17**. An overview of studies examining the feasibility of outplanting sexually reared *Acropora*, table from Guest et al. (2023) and supplemented with new studies. It shows the range of nursery time, outplant size, cost per nursery coral, survivorship, and cost per live coral from restoration ecology studies. Approximations are estimations from graphs and figures. When multiple outplant sites, species, size classes or devices were used, the best performing one was selected. When not reported, device yield was calculated by multiplying retention and survivorship, or when retention was not reported, survivorship was used as this value, unless devices were deployed in a way that allowed retrieval.

| *Reference* | *Species* | *Country* | *Age at outplant (mo)* | *Dia-meter at outplant (cm)* | *Cost of coral before outplant (US$)* | *Cost of outplanted coral (US$)* | *Age at final survey (mo)* | *Dia-meter final survey (cm)* | *Retention (%)* | *Device-level survi-vorship (%)* | *Device yield (ret * surv) (%)* | *Cost of outplant with live coral in study (US$)* | *Years old at costing* |
|---|---|---|---|---|---|---|---|---|---|---|---|---|---|
| 1. Omori et al. (2008) | *A. tenuis* | Japan | 18 | 5.8 | NA | NA | 24 | 9.1 | NA | 89 | 89 | NA | NA |
| 2. Boch and Morse (2012) | *A. hyacinthus* | Palau | 0.3 | ~0.1 | NA | NA | 12 | 1.3 | NA | 24.6 | 24.6 | NA | NA |
| 3. Villanueva et al. (2012) | *A. valida* | Philippines | 6 | 1.1 | 4.4 | 5.3 | 12 | 3.8 | 89.3 | 67.5 | 60.3 | 11.2 | 1 |
| 4. Guest et al. (2014) | *A. millepora* | Philippines | 7 | 0.5 | 19 | 23 | 30 | ~8.1 | NA | 8.3 | NA | 284 | 2.5 |
| | | | 14 | 2.3 | 22 | 26 | 30 | 7.6 | NA | 11.7 | NA | 217 | 2.5 |
| | | | 19 | 5.9 | 25 | 29 | 30 | 8.6 | NA | 46.7 | NA | 61 | 2.5 |
| 5. Chamberland et al. (2015) | *A. palmata* | Curaçao | 0.5 | 0.3 | 3.5 | 4.0 | 31 | ~4.6 | NA | 32 | NA | 12.6 | 2.5 |
| 6. Baria-Rodriguez et al. (2019) | *A. granulosa* | Philippines | 12 | 2.3 | 2.79 | NA | 25 | ~8.4 | NA | 18 | 18 | 20.01 | 2.1 |
| 7. Ligson et al. (2019) | *A. verweyi* | Philippines | 4 | 1.3 | 1.52 | 2.67 | 16 | ~3.9 | NA | 35.8 | NA | 11.47 | 1 |
| 8. Humanes et al. (2021) | *A. digitifera* | Palau | 5 | NA | 4.46 | 9.24 | 32 | ~1.1 | NA | 6 | NA | 227 | 2.5 |
| | | | 11 | NA | 5.60 | 10.39 | 32 | ~1.6 | NA | 30 | NA | 49 | 2.5 |
| 9. Randall et al. (2022) | *A. tenuis* | Australia | 3.5 | NA | NA | NA | 11 | NA | NA | 29 | NA | NA | NA |
| 10. Whitman et al. (2024) | *A. digitifera* | Australia | 5 | <0.8 | NA | NA | 13 | NA | NA | 23 | 23 | NA | NA |
| 11. Waters et al. (2025) | *A. spp* | Australia | 0.3 | ~0.1 | NA | NA | 21 | NA | NA | 9 | 9 | NA | NA |
| 12. Whitman et al. (2025) | *A. digitifera* | Australia | 2 | 0.2 | NA | NA | 18 | 1.4 | NA | 57 | 57 | NA | NA |
| 13. van der Steeg et al. (2025) | *A. digitifera* | Palau | 6 | 2.4 | NA | NA | 20 | 3.8 | NA | 78 | 65 | NA | NA |
| 14. Ramsby et al. (2025) | *A. spathulate* | Australia | ~1 | ~0.1 | NA | NA | 13 | NA | 89 | 60 | 53.4 | NA | NA |
| | *A. digitifera* | | ~2 | ~0.2 | NA | NA | 16 | NA | 62 | 24 | 14.9 | NA | NA |
| | *A. spathulate* | | ~0.5 | ~0.1 | NA | NA | 5 | NA | 65 | 87 | 56.6 | NA | NA |
| | *A. millepora* | | ~0.5 | ~0.1 | NA | NA | 8 | NA | 61 | 73 | 44.5 | NA | NA |
| 15. CoralAssist Plug | *A. digitifera* | Palau | 1 | ~0.1 | 8.98 | 13.97 | 38 | 5.6 | 90.7 | 25.0 | 24.0 | 79.50 | 3 |
| | | | 3 | ~0.4 | 13.37 | 19.09 | 50 | 10.5 | 62.0 | 32.4 | 16.0 | 168.12 | 4 |
| | | | 4 | ~0.6 | NA | NA | 38 | 6.3 | 84.6 | 48.6 | 36.5 | NA | 3 |
| | | | 6 | ~1.5 | 16.25 | 22.92 | 50 | 9.4 | 70.0 | 52.3 | 36.0 | 89.00 | 4 |
