## Appendix S2 for "The CoralAssist Plug: a novel device that enhances outplant rate and survivorship of sexually propagated corals"

### Table S1. Experimental Costs

| **Local salary rates** | Annual | Monthly | **Hourly rate** |
|---|---|---|---|
| Research assistant | $12,000 | $1,000 | $6.25 |
| Scientific expert | $40,000 | $3,333 | $20.83 |

| Local rates of scuba tank rental and boat hire | | | |
|---|---|---|---|
| Boat full day | $375 | Scuba tank | $7 |
| Boat half day | $220 | | |
| Fuel per day | $25 | | |

| | | Material cost | | | Personnel cost | | | Boat & fuel | | SCUBA tanks | | Total cost/yr | % total |
|---|---|---|---|---|---|---|---|---|---|---|---|---|---|
| | **Activity** | Capital | Consumables | USD | Person hours | Rate | USD | Boat days | USD | # tanks | USD | USD | cost/yr |
| 1 | Collection adult colonies | $1,673 | $16 | $1,689 | 16 | $13.54 | $217 | 2 half | $490 | 8 | $56 | $1,336 | 7% |
| 2 | Spawning & larvae rearing | $1,674 | $44 | $1,718 | 98 | $11.90 | $1,167 | 0 | $0 | 0 | $0 | $1,769 | 9% |
| 3 | Substrates & Conditioning | $3,194 | $2,162 | $5,356 | 16 | $11.90 | $184 | 1 half | $245 | 4 | $28 | $3,684 | 20% |
| 4 | Flow through settlement tanks 1-month | $129 | $16 | $145 | 17 | $6.25 | $106 | 0 | $0 | 0 | $0 | $165 | 1% |
| 5 | Ex-situ nursery 3-months | $4,879 | $232 | $5,111 | 60 | $6.25 | $375 | 0 | $0 | 0 | $0 | $2,234 | 12% |
| 5.6 | Ex-situ nursery 6-months, additional | $0 | $76 | $76 | 48 | $6.25 | $300 | 0 | $0 | 0 | $0 | $376 | 2% |
| 6 | Outplanting (n=600) | $503 | $1,541 | $2,044 | 69 | $11.96 | $825 | 3 full | $1,200 | 18 | $126 | $3,860 | 21% |
| 7 | Monitoring after 1 year (n=600, 1.5 days) | $1,158 | $59 | $1,218 | 128 | $15.36 | $1,967 | 4 full, 4 half | $2,580 | 40 | $280 | $5,272 | 28% |
| | SUM | $13,210 | $4,147 | $17,357 | 452 | | $5,141 | 7 full, 7 half | $4,515 | 70 | $490 | $18,696 | 100% |

| Data this study | | |
|---|---|---|
| Eggs collected from spawning | 872,500 | 3 colonies, other 7 spawned day after; 766700 eggs |
| Fertilization | 94.6% | in 2021 ~94% |
| Larvae survivorship 4 days | 77.0% | in 2021 ~27% |
| Larvae settlement success | 44.1% | in 2021 22.4% |
| Max substrates for settlemet at 50 larvae/sub | 12,711 | |

| | *CAP* | *Control* | *Combined* | |
|---|---|---|---|---|
| Number of substrates settled | 450 | 450 | 900 | |
| Percentage successful settled | 87.5% | 81.3% | 84.4% | 2020 subset |
| Nursery yield after 1 month | 86.0% | 86.0% | 86.0% | 2021 full count |
| Nursery yield after 3 months | 75.0% | 62.5% | 68.8% | From 2020 subset |
| Nursery yield after 4 months | 78.9% | 83.0% | 81.1% | 2021 full count |
| Nursery yield after 6 months | 64.3% | 48.1% | 56.1% | Not available, assumed same as 3m nursery yield, but likely better |

### Table S1. Experimental Costs

| 2020 Spawning | *CAP* | *CAP* | *Control* | *Control* | *Combined* | *Combined* |
|---|---|---|---|---|---|---|
| Outplant age | 3 months | 6 months | 3 months | 6 months | 3 months | 6 months |
| Outplant yield after 1 year | 57% | 67% | 40% | 49% | 49% | 58% |
| Outplant yield after 2 years | 45% | 60% | 28% | 38% | 37% | 49% |
| Outplant yield after 3 years | 26% | 47% | 20% | 31% | 23% | 39% |
| Outplant yield after 4 years | 16% | 36% | 17% | 25% | 17% | 31% |

| 2021 Spawning | *CAP* | *CAP* | *Control* | *Control* | *Combined* | *Combined* | |
|---|---|---|---|---|---|---|---|
| Outplant age | 1 month | 4 months | 1 month | 4 months | 1 month | 4 months | 4 month had combined flow-through and normal |
| Outplant yield after 1 year | 53% | 59% | 45% | 54% | 49% | 56% | nursery so would be very expensive |
| Outplant yield after 2 years | 33% | 45% | 31% | 36% | 32% | 41% | |
| Outplant yield after 3 years | 24% | 37% | 17% | 23% | 21% | 30% | |

| | | | | |
|---|---|---|---|---|
| Device yields in nurseries | 900 | | | |
| Settled | 760 | | | |
| 1-month-old corals | 774 | 574 | | 200 outplanted at 1m |
| 3-month-old corals | 619 | 468 | 268 | 200 outplanted at 3m |
| 6-month-old corals | 505 | 381 | 218 | 200 outplanted at 6m |

#### Table S1. Experimental Costs

| Cost per live coral | *CAP* | *Control* | *Combined* |
|---|---|---|---|
| 1-month-old in flow-through tanks | $8.98 | $8.98 | $8.98 |
| 3-month-old in nursery | $13.37 | $16.03 | $14.57 |
| 6-month-old in nursery | $16.25 | $21.72 | $18.63 |
| 1-month-old outplant | $13.97 | $13.97 | $13.97 |
| 3-month-old outplant | $19.09 | $22.89 | $20.81 |
| 6-month-old outplant | $22.92 | $30.64 | $26.28 |
| 1-year-old outplanted at 1-month-old | $29.41 | $34.60 | $31.79 |
| 1-year-old outplanted at 3-months-old | $36.91 | $63.09 | $47.29 |
| 1-year-old outplanted at 6-months-old | $37.62 | $68.75 | $49.81 |
| 2-year-old outplanted at 1-months-old | $52.18 | $56.60 | $54.30 |
| 2-year-old outplanted at 3-months-old | $51.10 | $98.50 | $68.68 |
| 2-year-old outplanted at 6-months-old | $45.80 | $96.67 | $64.29 |
| 3-year-old outplanted at 1-month-old | $79.50 | $110.29 | $92.17 |
| 3-year-old outplanted at 3-months-old | $95.95 | $149.61 | $118.24 |
| 3-year-old outplanted at 6-months-old | $63.32 | $128.32 | $87.47 |
| 4-year-old outplanted at 3-months-old | $168.12 | $189.79 | $177.73 |
| 4-year-old outplanted at 6-months-old | $89.00 | $171.30 | $120.42 |

Cost of CAP and control devided over 2 as those are spread over the two devices

Devide monitoring costs per years, multiply with outplant yield after x years

### Table S2. Realistic Rehabilitation Costs

| Local salary rates | Annual | Monthly | Hourly rate |
|---|---|---|---|
| Research assistant | $12,000 | $1,000 | $6.25 |
| Scientific expert | $40,000 | $3,333 | $20.83 |

| Local rates of scuba tank rental and boat hire | | | |
|---|---|---|---|
| Boat full day | $375 | Scuba tank | $7 |
| Boat half day | $220 | | |
| Fuel per day | $25 | | |

| | | Material cost | | | Personnel cost | | | Boat & fuel | | SCUBA tanks | |
|---|---|---|---|---|---|---|---|---|---|---|---|
| | **Activity** | Capital | Consumables | USD | Person hours | Rate | USD | Boat days | USD | # tanks | USD |
| 1 | Collection adult colonies | $1,610 | $16 | $1,626 | 16 | $13.54 | $217 | 2 half | $490 | 8 | $56 |
| 2 | Spawning & larvae rearing | $2,593 | $45 | $2,638 | 162 | $9.67 | $1,567 | 0 | $0 | 0 | $0 |
| 3 | Substrates & Conditioning | $745 | $10,600 | $11,345 | 40 | $8.44 | $338 | 1 half | $245 | 4 | $28 |
| 4 | Flow through settlement tanks 1-month | $839 | $124 | $964 | 68 | $6.25 | $425 | 0 | $0 | 0 | $0 |
| 5 | Ex-situ nursery 3-months | $13,866 | $1,051 | $14,916 | 200 | $6.25 | $1,250 | 0 | $0 | 0 | $0 |
| 5.6 | Ex-situ nursery 6-months, additional | $0 | $437 | $437 | 192 | $6.25 | $1,200 | 0 | $0 | 0 | $0 |
| 6 | Outplanting | $5,396 | $2,869 | $8,265 | 0 | 0 | $0 | 0 | $0 | 0 | $0 |
| 6.1 | Outplanting all at 1-month-old | $0 | $573 | $573 | 671 | $8.60 | $5,769 | 13 full | $5,200 | 234 | $1,638 |
| 6.3 | Outplanting all at 3-months-old | $0 | $465 | $465 | 569 | $8.61 | $4,898 | 11 full | $4,400 | 198 | $1,386 |
| 6.6 | Outplanting all at 6-months-old | $0 | $384 | $384 | 467 | $8.62 | $4,027 | 9 full | $3,600 | 162 | $1,134 |
| 7 | Monitoring subset yearly (n=400, 4 days) | $490 | $66 | $557 | 96 | $15.97 | $1,533 | 4 full | $1,600 | 24 | $168 |
| | SUM | $25,539 | $16,631 | $42,169 | 2385 | | $21,223 | 4 full, 4 half | $15,535 | 630 | $4,410 |

#### Table S2. Realistic Rehabilitation Costs

Table continued; Total costs

| | | 1-month-old outplant | | 3-month-old outplant | | 6-month-old outplant | |
|---|---|---|---|---|---|---|---|
| | | Total cost/yr | % total | Total cost/yr | % total | Total cost/yr | % total |
| | **Activity** | USD | cost/yr | USD | cost/yr | USD | cost/yr |
| 1 | Collection adult colonies | $1,315 | 4% | $1,315 | 3% | $1,315 | 3% |
| 2 | Spawning & larvae rearing | $2,476 | 7% | $2,476 | 6% | $2,476 | 6% |
| 3 | Substrates & Conditioning | $11,459 | 31% | $11,459 | 28% | $11,459 | 28% |
| 4 | Flow through settlement tanks 1-month | $829 | 2% | $0 | 0% | $0 | 0% |
| 5 | Ex-situ nursery 3-months | $0 | 0% | $6,923 | 17% | $6,923 | 17% |
| 5.6 | Ex-situ nursery 6-months, additional | $0 | 0% | $0 | 0% | $1,637 | 4% |
| 6 | Outplanting | $4,668 | 12% | $4,668 | 11% | $4,668 | 11% |
| 6.1 | Outplanting all at 1-month-old | $13,180 | 35% | $0 | 0% | $0 | 0% |
| 6.3 | Outplanting all at 3-months-old | $0 | 0% | $11,149 | 27% | $0 | 0% |
| 6.6 | Outplanting all at 6-months-old | $0 | 0% | $0 | 0% | $9,145 | 22% |
| 7 | Monitoring subset yearly (n=400, 4 days) | $3,531 | 9% | $3,531 | 9% | $3,531 | 9% |
| | SUM | $37,457 | 100% | $41,520 | 100% | $41,154 | 100% |

| Data this study | | |
|---|---|---|
| Eggs collected from spawning | 1,639,200 | 3 colonies, other 7 spawned day after; 766700 eggs, total= 1,639,200, could settle 23,880 substrates |
| Fertilization | 94.6% | in 2021 ~94% |
| Larvae survivorship 4 days | 77.0% | in 2021 ~27% |
| Larvae settlement success | 44.1% | in 2021 22.4% |
| Max substrates for settlemet at 50 larvae/sub | 23,881 | |

| | *CAP* | |
|---|---|---|
| Number of substrates settled | 10,000 | |
| Percentage successful settled | 87.5% | 2020 subset |
| Nursery yield after 1 month | 86.0% | 2021 full count |
| Nursery yield after 3 months | 75.0% | 2020 subset 0.3-3m yield 85.7%, in 2021 1-4m yield was: 94.3% |
| Nursery yield after 6 months | 64.3% | Not available, assumed same as 3m nursery yield, but likely better |

### Table S2. Realistic Rehabilitation Costs

| 2020 Spawning | *CAP* | *CAP* |
|---|---|---|
| Outplant age | 3 months | 6 months |
| Outplant yield after 1 year | 57% | 67% |
| Outplant yield after 2 years | 45% | 60% |
| Outplant yield after 3 years | 26% | 47% |
| Outplant yield after 4 years | 16% | 36% |

| 2021 Spawning | *CAP* | *CAP* |
|---|---|---|
| Outplant age | 1 month | 4 months |
| Outplant yield after 1 year | 53% | 59% |
| Outplant yield after 2 years | 33% | 45% |
| Outplant yield after 3 years | 24% | 37% |

| | | | |
|---|---|---|---|
| Substrate yields in nurseries | 10000 | | |
| Settled | 8750 | 700 outplants per day with 2 teams of 3 divers | |
| 1-month-old corals | 8600 | 12 | Days needed for outplant |
| 3-month-old corals | 7499 | 11 | |
| 6-month-old corals | 6426 | 9 | |

| Cost per live coral | *CAP* |
|---|---|
| 1-month-old in flow-through tanks | $1.87 |
| 3-month-old in nursery | $2.96 |
| 6-month-old in nursery | $3.70 |
| 1-month-old outplant | $3.94 |
| 3-month-old outplant | $5.07 |
| 6-month-old outplant | $5.85 |
| 1-year-old outplanted at 1-month-old | $7.59 |
| 1-year-old outplanted at 3-months-old | $9.09 |
| 1-year-old outplanted at 6-months-old | $8.94 |
| 2-year-old outplanted at 1-month-old | $12.46 |
| 2-year-old outplanted at 3-months-old | $11.78 |
| 2-year-old outplanted at 6-months-old | $10.22 |
| 3-year-old outplanted at 1-month-old | $17.72 |
| 3-year-old outplanted at 3-months-old | $20.84 |
| 3-year-old outplanted at 6-months-old | $13.33 |
| 4-year-old outplanted at 1-month-old | NA |
| 4-year-old outplanted at 3-months-old | $34.61 |
| 4-year-old outplanted at 6-months-old | $17.79 |

#### Table S3. Materials

| Act | Activity | Item | Equipment/ Consumables | Quantity exp. | Cost $ experiment | Total experiment | Notes exp. | Quantity scaled | Cost $ scaled | Total scaled | Notes scaled |
|---|---|---|---|---|---|---|---|---|---|---|---|
| 1 | Collection adult colonies | Dive gear | Equipment | 2 | $ 1,000.00 | $ 285.71 | Pro rata over 14 days | 2 | $ 1,000.00 | $ 222.22 | Pro rata over 18 days (average of 11 outplant days) |
| 1 | Collection adult colonies | Steel sledge hammer | Equipment | 1 | $ 28.99 | $ 28.99 | | 1 | $ 28.99 | $ 28.99 | |
| 1 | Collection adult colonies | Chisel with target guard | Equipment | 1 | $ 13.99 | $ 13.99 | | 1 | $ 13.99 | $ 13.99 | |
| 1 | Collection adult colonies | Cooler/tanks for transport | Equipment | 3 | $ 24.95 | $ 74.85 | | 3 | $ 24.95 | $ 74.85 | |
| 1 | Collection adult colonies | Holding tank 400L | Equipment | 1 | $ 900.00 | $ 900.00 | | 1 | $ 900.00 | $ 900.00 | |
| 1 | Collection adult colonies | Magnetic drive pump | Equipment | 3 | $ 119.99 | $ 359.97 | | 3 | $ 119.99 | $ 359.97 | |
| 1 | Collection adult colonies | Coral stands 4" PVC (per foot) | Equipment | 2 | $ 4.76 | $ 9.52 | | 2 | $ 4.76 | $ 9.52 | |
| 1 | Collection adult colonies | Filter sock 50µm large | Consumable | 2 | $ 8.00 | $ 16.00 | | 2 | $ 8.00 | $ 16.00 | |
| 2 | Spawning & larvae rearing | Red headlamp | Equipment | 2 | $ 19.99 | $ 39.98 | | 2 | $ 19.99 | $ 39.98 | |
| 2 | Spawning & larvae rearing | Collection cups | Equipment | 4 | $ 1.20 | $ 4.80 | | 4 | $ 1.20 | $ 4.80 | |
| 2 | Spawning & larvae rearing | Pentek filter vessel | Equipment | 1 | $ 289.88 | $ 289.88 | | 1 | $ 289.88 | $ 289.88 | |
| 2 | Spawning & larvae rearing | Corner brace filter vessel | Equipment | 1 | $ 5.59 | $ 5.59 | | 1 | $ 5.59 | $ 5.59 | |
| 2 | Spawning & larvae rearing | 3 compartment filter | Equipment | 1 | $ 317.06 | $ 317.06 | | 1 | $ 317.06 | $ 317.06 | |
| 2 | Spawning & larvae rearing | UV filter | Equipment | 1 | $ 465.85 | $ 465.85 | | 1 | $ 465.85 | $ 465.85 | |
| 2 | Spawning & larvae rearing | PVC pipe 1" (foot) | Equipment | 6 | $ 0.60 | $ 3.59 | | 6 | $ 0.60 | $ 3.59 | |
| 2 | Spawning & larvae rearing | PVC valve 1" | Equipment | 1 | $ 2.95 | $ 2.95 | | 1 | $ 2.95 | $ 2.95 | |
| 2 | Spawning & larvae rearing | PVC elbow 1" | Equipment | 1 | $ 0.85 | $ 0.85 | | 1 | $ 0.85 | $ 0.85 | |
| 2 | Spawning & larvae rearing | PVC union 1" | Equipment | 1 | $ 4.65 | $ 4.65 | | 1 | $ 4.65 | $ 4.65 | |
| 2 | Spawning & larvae rearing | PVC adapter treaded male 1" | Equipment | 2 | $ 1.29 | $ 2.58 | | 2 | $ 1.29 | $ 2.58 | |
| 2 | Spawning & larvae rearing | PVC adapter 1" to 3/4" threaded | Equipment | 1 | $ 0.85 | $ 0.85 | | 1 | $ 0.85 | $ 0.85 | |
| 2 | Spawning & larvae rearing | PVC union 3/4" | Equipment | 2 | $ 4.49 | $ 8.98 | | 2 | $ 4.49 | $ 8.98 | |
| 2 | Spawning & larvae rearing | PVC pipe 3/4" (foot) | Equipment | 1 | $ 0.44 | $ 0.44 | | 1 | $ 0.44 | $ 0.44 | |
| 2 | Spawning & larvae rearing | PVC adapter 3/4" to 1/2" threaded | Equipment | 1 | $ 0.85 | $ 0.85 | | 1 | $ 0.85 | $ 0.85 | |
| 2 | Spawning & larvae rearing | PVC pipe 1/2" | Equipment | 6 | $ 0.31 | $ 1.88 | | 6 | $ 0.31 | $ 1.88 | |
| 2 | Spawning & larvae rearing | PVC valve 1/2" | Equipment | 1 | $ 1.55 | $ 1.55 | | 1 | $ 1.55 | $ 1.55 | |
| 2 | Spawning & larvae rearing | PVC tee threaded 1/2" | Equipment | 1 | $ 0.55 | $ 0.55 | | 1 | $ 0.55 | $ 0.55 | |
| 2 | Spawning & larvae rearing | PVC stop 1/2" | Equipment | 1 | $ 0.50 | $ 0.50 | | 1 | $ 0.50 | $ 0.50 | |
| 2 | Spawning & larvae rearing | Rearing tanks | Equipment | 8 | $ 17.00 | $ 136.00 | | 34 | $ 17.00 | $ 578.00 | |
| 2 | Spawning & larvae rearing | Banjo filter 4" PVC (foot) | Equipment | 1 | $ 4.76 | $ 4.76 | | 2 | $ 4.76 | $ 9.52 | |
| 2 | Spawning & larvae rearing | PVC pipe 3/4" (foot) | Equipment | 0.5 | $ 0.44 | $ 0.22 | | 1 | $ 0.44 | $ 0.44 | |
| 2 | Spawning & larvae rearing | PVC pipe cutter | Equipment | 1 | $ 15.55 | $ 15.55 | | 1 | $ 15.55 | $ 15.55 | |
| 2 | Spawning & larvae rearing | PVC connector 3/4" threaded f | Equipment | 3 | $ 1.29 | $ 3.87 | | 6 | $ 1.29 | $ 7.74 | |

#### Table S3. Materials

| Act | Activity | Item | Equipment/ Consumables | Quantity exp. | Cost $ experiment | Total experiment | Notes exp. | Quantity scaled | Cost $ scaled | Total scaled | Notes scaled |
|---|---|---|---|---|---|---|---|---|---|---|---|
| 2 | Spawning & larvae rearing | Plankton mesh 100µm (m2) | Equipment | 0.5 | $ 31.06 | $ 15.53 | | 1 | $ 31.06 | $ 31.06 | |
| 2 | Spawning & larvae rearing | Hose connector 3/4" threaded m | Equipment | 3 | $ 2.39 | $ 7.17 | | 6 | $ 2.39 | $ 14.34 | |
| 2 | Spawning & larvae rearing | Hose 1/2" threaded (foot) | Equipment | 20 | $ 0.95 | $ 19.00 | | 40 | $ 0.95 | $ 38.00 | |
| 2 | Spawning & larvae rearing | Drill core 3/4" | Equipment | 1 | $ 7.85 | $ 7.85 | | 1 | $ 7.85 | $ 7.85 | |
| 2 | Spawning & larvae rearing | Airpump+C24 | Equipment | 1 | $ 256.00 | $ 256.00 | | 2 | $ 256.00 | $ 512.00 | |
| 2 | Spawning & larvae rearing | Airline (m) | Equipment | 30 | $ 1.17 | $ 35.10 | | 120 | $ 1.17 | $ 140.40 | |
| 2 | Spawning & larvae rearing | Airline adjusters | Equipment | 8 | $ 1.29 | $ 10.32 | | 34 | $ 1.29 | $ 43.86 | |
| 2 | Spawning & larvae rearing | Rigid airline | Equipment | 8 | $ 1.20 | $ 9.60 | | 34 | $ 1.20 | $ 40.80 | |
| 2 | Spawning & larvae rearing | Filter sock 50µm small | Consumable | 2 | $ 8.00 | $ 16.00 | | 2 | $ 8.00 | $ 16.00 | |
| 2 | Spawning & larvae rearing | Filter catridge 10µm | Consumable | 1 | $ 2.15 | $ 2.15 | | 1 | $ 2.15 | $ 2.15 | |
| 2 | Spawning & larvae rearing | Filter catridge 5µm | Consumable | 2 | $ 2.85 | $ 5.70 | | 2 | $ 2.85 | $ 5.70 | |
| 2 | Spawning & larvae rearing | Filter catridge 1µm | Consumable | 2 | $ 2.59 | $ 5.18 | | 2 | $ 2.59 | $ 5.18 | |
| 2 | Spawning & larvae rearing | Super glue | Consumable | 1 | $ 4.84 | $ 4.84 | | 1 | $ 4.84 | $ 4.84 | |
| 2 | Spawning & larvae rearing | Cable ties | Consumable | 100 | $ 0.01 | $ 1.00 | | 200 | $ 0.01 | $ 2.00 | |
| 2 | Spawning & larvae rearing | PVC glue | Consumable | 1 | $ 8.95 | $ 8.95 | | 1 | $ 8.95 | $ 8.95 | |
| 3 | Substrates & Conditioning | Dive gear | Equipment | 2 | $ 1,000.00 | $ 142.86 | Pro rata over 14 days | 2 | $ 1,000.00 | $ 111.11 | Pro rata over 18 days |
| 3 | Substrates & Conditioning | Mesh catch bag | Equipment | 1 | $ 21.00 | $ 21.00 | | 1 | $ 21.00 | $ 21.00 | |
| 3 | Substrates & Conditioning | Garden stake 2ft | Equipment | 20 | $ 1.70 | $ 34.00 | | 200 | $ 1.70 | $ 340.00 | |
| 3 | Substrates & Conditioning | PVC pipe 2" (foot) | Equipment | 10 | $ 1.60 | $ 16.00 | | 100 | $ 1.60 | $ 160.00 | |
| 3 | Substrates & Conditioning | Shading cloth | Equipment | 1 | $ 12.94 | $ 12.94 | | 4 | $ 12.94 | $ 51.76 | |
| 3 | Substrates & Conditioning | PVC Pipe 1/2" | Equipment | 36 | $ 0.31 | $ 11.16 | | 144 | $ 0.31 | $ 44.64 | |
| 3 | Substrates & Conditioning | PVC Elbow 1/2" | Equipment | 8 | $ 0.50 | $ 4.00 | | 32 | $ 0.50 | $ 16.00 | |
| 3 | Substrates & Conditioning | Production tools CAPs & controls | Equipment | 2 | $ 1,476.00 | $ 2,952.00 | | 0 | $ - | $ - | |
| 3 | Substrates & Conditioning | CAPs and control devices | Consumable | 900 | $ 2.18 | $ 1,962.00 | | 10000 | $ 1.01 | $ 10,100.00 | New production method |
| 3 | Substrates & Conditioning | Shipping | Consumable | 1 | $ 200.00 | $ 200.00 | | 1 | $ 500.00 | $ 500.00 | |
| 4 | Flow through 1m outplant | Plastic container 20L | Equipment | 3 | $ 11.95 | $ 35.85 | | 0 | $ 11.95 | $ - | Use larvae rearing containers |
| 4 | Flow through 1m outplant | Bulkhead fittings 3/4 to 1/2" | Equipment | 3 | $ 4.00 | $ 12.00 | | 34 | $ 4.00 | $ 136.00 | |
| 4 | Flow through 1m outplant | PVC pipe 3/4" (foot) | Equipment | 1 | $ 0.44 | $ 0.44 | | 11 | $ 0.44 | $ 4.84 | |
| 4 | Flow through 1m outplant | PVC Elbow 3/4" | Equipment | 3 | $ 0.70 | $ 2.10 | | 34 | $ 0.70 | $ 23.80 | |

#### Table S3. Materials

| Act | Activity | Item | Equipment/ Consumables | Quantity exp. | Cost $ experiment | Total experiment | Notes exp. | Quantity scaled | Cost $ scaled | Total scaled | Notes scaled |
|---|---|---|---|---|---|---|---|---|---|---|---|
| 4 | Flow through 1m outplant | Banjo filter 4" PVC (foot) | Equipment | 1 | $ 4.76 | $ 4.76 | | 4 | $ 4.76 | $ 19.04 | |
| 4 | Flow through 1m outplant | PVC pipe 3/4" (foot) | Equipment | 0.5 | $ 0.44 | $ 0.22 | | 2 | $ 0.44 | $ 0.88 | |
| 4 | Flow through 1m outplant | Plankton mesh 100µm (m2) | Equipment | 0.5 | $ 31.06 | $ 15.53 | | 2 | $ 31.06 | $ 62.12 | |
| 4 | Flow through 1m outplant | PVC tee threaded 1/2" | Equipment | 3 | $ 0.55 | $ 1.65 | | 34 | $ 0.55 | $ 18.70 | |
| 4 | Flow through 1m outplant | John Guest 1/2 to 1/4 connector | Equipment | 6 | $ 3.68 | $ 22.08 | | 68 | $ 3.68 | $ 250.24 | |
| 4 | Flow through 1m outplant | John Guest valve adjusters | Equipment | 3 | $ 8.00 | $ 24.00 | | 34 | $ 8.00 | $ 272.00 | |
| 4 | Flow through 1m outplant | Water tube 1/4" (meter) | Equipment | 8 | $ 1.29 | $ 10.32 | | 40 | $ 1.29 | $ 51.60 | |
| 4 | Flow through 1m outplant | Super glue | Consumable | 1 | $ 4.84 | $ 4.84 | | 3 | $ 4.84 | $ 14.52 | |
| 4 | Flow through 1m outplant | Coral food | Consumable | 0.5 | $ 21.98 | $ 10.99 | | 5 | $ 21.98 | $ 109.90 | |
| 5 | Ex-situ nursery | Marine wood for table 4x8 ft 1/2" | Equipment | 2 | $ 49.45 | $ 98.90 | | 0 | $ 49.45 | $ - | |
| 5 | Ex-situ nursery | Wood 2x4 (foot) | Equipment | 216 | $ 1.00 | $ 215.40 | | 0 | $ 1.00 | $ - | |
| 5 | Ex-situ nursery | Screws 10" (pound) | Equipment | 2 | $ 2.95 | $ 5.90 | | 0 | $ 2.95 | $ - | |
| 5 | Ex-situ nursery | Nursery tank 184L | Equipment | 4 | $ 400.00 | $ 1,600.00 | | 0 | $ 400.00 | $ - | |
| 5 | Ex-situ nursery | Pentek filter vessel | Equipment | 0 | $ 289.88 | $ - | From spawning | 0 | $ 289.88 | $ - | |
| 5 | Ex-situ nursery | Corner brace filter vessel | Equipment | 0 | $ 5.59 | $ - | From spawning | 0 | $ 5.59 | $ - | |
| 5 | Ex-situ nursery | PVC pipe 1" (foot) | Equipment | 74 | $ 0.60 | $ 44.22 | | 0 | $ 0.60 | $ - | |
| 5 | Ex-situ nursery | PVC union 1" | Equipment | 2 | $ 4.65 | $ 9.30 | | 0 | $ 4.65 | $ - | |
| 5 | Ex-situ nursery | PVC elbow 1" | Equipment | 27 | $ 0.85 | $ 22.95 | | 0 | $ 0.85 | $ - | |
| 5 | Ex-situ nursery | PVC tee 1" | Equipment | 8 | $ 1.56 | $ 12.48 | | 0 | $ 1.56 | $ - | |
| 5 | Ex-situ nursery | PVC valve 1" | Equipment | 6 | $ 2.95 | $ 17.70 | | 0 | $ 2.95 | $ - | |
| 5 | Ex-situ nursery | PVC adapter treaded male 1" | Equipment | 4 | $ 1.29 | $ 5.16 | | 0 | $ 1.29 | $ - | |
| 5 | Ex-situ nursery | PVC adapter treaded female 1" | Equipment | 4 | $ 1.29 | $ 5.16 | | 0 | $ 1.29 | $ - | |
| 5 | Ex-situ nursery | Powerstrip 4 normal & 4 timed outlets | Equipment | 2 | $ 22.99 | $ 45.98 | | 4 | $ 22.99 | $ 91.96 | |
| 5 | Ex-situ nursery | Pumps Koralia Evolution 850 | Equipment | 8 | $ 39.99 | $ 319.92 | | 32 | $ 39.99 | $ 1,279.68 | |
| 5 | Ex-situ nursery | Lights LED | Equipment | 8 | $ 296.99 | $ 2,375.92 | | 32 | $ 296.99 | $ 9,503.68 | |
| 5 | Ex-situ nursery | Holding drain for lights (foot) | Equipment | 36 | $ 1.00 | $ 36.00 | | 144 | $ 1.00 | $ 144.00 | |
| 5 | Ex-situ nursery | Drain covers | Equipment | 16 | $ 3.99 | $ 63.84 | | 32 | $ 3.99 | $ 127.68 | |
| 5 | Ex-situ nursery | Holding tank 400L | Equipment | 0 | $ 900.00 | $ - | | 3 | $ 900.00 | $ 2,700.00 | Use 3 additional large holding tanks for rearing |

#### Table S3. Materials

| Act | Activity | Item | Equipment/ Consumables | Quantity exp. | Cost $ experiment | Total experiment | Notes exp. | Quantity scaled | Cost $ scaled | Total scaled | Notes scaled |
|---|---|---|---|---|---|---|---|---|---|---|---|
| 5 | Ex-situ nursery | PVC Pipe 1/2" (foot) frame substrates | Equipment | 0 | $ 0.31 | $ - | | 30 | $ 0.31 | $ 9.30 | |
| 5 | Ex-situ nursery | PVC side outlet elbow 1/2" | Equipment | 0 | $ 1.55 | $ - | | 4 | $ 1.55 | $ 6.20 | |
| 5 | Ex-situ nursery | PVC tee 1/2" | Equipment | 0 | $ 0.55 | $ - | | 6 | $ 0.55 | $ 3.30 | |
| 5 | Ex-situ nursery | Filter sock 50µm small | Consumable | 2 | $ 5.00 | $ 10.00 | | 8 | $ 5.00 | $ 40.00 | |
| 5 | Ex-situ nursery | Rope for suspension 3/16" (foot) | Consumable | 64 | $ 0.22 | $ 14.07 | | 336 | $ 0.22 | $ 73.85 | |
| 5 | Ex-situ nursery | Pulley | Consumable | 16 | $ 3.99 | $ 63.84 | | 64 | $ 3.99 | $ 255.36 | |
| 5 | Ex-situ nursery | Cleat | Consumable | 16 | $ 2.95 | $ 47.20 | | 64 | $ 2.95 | $ 188.80 | |
| 5 | Ex-situ nursery | Coculturing rabbitfish | Consumable | 40 | $ - | $ - | Donated | 80 | $ - | $ - | Donated |
| 5 | Ex-situ nursery | Coculturing juvenile trochus snails | Consumable | 400 | $ - | $ - | Reared at PICRC | 800 | $ - | $ - | Reared at PICRC |
| 5 | Ex-situ nursery | Coculturing cerith snails | Consumable | 400 | $ - | $ - | Collected around PICRC | 800 | $ - | $ - | Collected around PICRC |
| 5 | Ex-situ nursery | Coculturing filefish | Consumable | 4 | $ - | $ - | Collected around PICRC | 4 | $ - | $ - | Collected around PICRC |
| 5 | Ex-situ nursery | Fish food | Consumable | 2 | $ 27.16 | $ 54.32 | | 8 | $ 27.16 | $ 217.28 | |
| 5 | Ex-situ nursery | Coral food | Consumable | 1 | $ 21.98 | $ 21.98 | | 10 | $ 21.98 | $ 219.80 | |
| 5 | Ex-situ nursery | Brushes | Consumable | 2 | $ 4.62 | $ 9.24 | | 2 | $ 4.62 | $ 9.24 | |
| 5 | Ex-situ nursery | Bleach | Consumable | 1 | $ 4.99 | $ 4.99 | | 4 | $ 4.99 | $ 19.96 | |
| 5 | Ex-situ nursery | Vinegar | Consumable | 1 | $ 6.59 | $ 6.59 | | 4 | $ 6.59 | $ 26.36 | |
| 5.6 | Ex-situ nursery | Fish food | Consumable | 2 | $ 27.16 | $ 54.32 | | 8 | $ 27.16 | $ 217.28 | |
| 5.6 | Ex-situ nursery | Coral food | Consumable | 1 | $ 21.98 | $ 21.98 | | 10 | $ 21.98 | $ 219.80 | |
| 6 | Outplanting | Dive gear | Equipment | 2 | $ 1,000.00 | $ 428.57 | Pro rata over 14 days | 6 | $ 1,000.00 | $ 5,222.22 | Pro rata over 18 & 11 days |
| 6 | Outplanting | Transect tape 50m | Equipment | 1 | $ 40.58 | $ 40.58 | | 2 | $ 40.58 | $ 81.16 | |
| 6 | Outplanting | Plastic basket | Equipment | 1 | $ 2.25 | $ 2.25 | | 4 | $ 2.25 | $ 9.00 | |
| 6 | Outplanting | 1 lb dive weight | Equipment | 1 | $ 10.50 | $ 10.50 | | 4 | $ 10.50 | $ 42.00 | |
| 6 | Outplanting | Mesh catch bag | Equipment | 1 | $ 21.00 | $ 21.00 | | 2 | $ 21.00 | $ 42.00 | |
| 6 | Outplanting | Nemo underwater drill | Consumable | 1 | $ 1,386.00 | $ 1,386.00 | | 2 | $ 1,386.00 | $ 2,772.00 | |
| 6 | Outplanting | Drill bit 5/16" | Consumable | 3 | $ 10.99 | $ 32.97 | | 4 | $ 10.99 | $ 43.96 | |
| 6 | Outplanting | 1ft nails (2 per transect + spare) | Consumable | 28 | $ 2.49 | $ 69.72 | 12 transects | 18 | $ 2.49 | $ 44.82 | 8 transects for subset monitoring |

#### Table S3. Materials

| Act | Activity | Item | Equipment/ Consumables | Quantity exp. | Cost $ experiment | Total experiment | Notes exp. | Quantity scaled | Cost $ scaled | Total scaled | Notes scaled |
|---|---|---|---|---|---|---|---|---|---|---|---|
| 6 | Outplanting | Cattle tags (2 per transect) | Consumable | 24 | $ 0.26 | $ 6.24 | | 16 | $ 0.26 | $ 4.16 | 8 transects for subset monitoring |
| 6 | Outplanting | Masonry nails 1-1/2" (pounds) | Consumable | 2 | $ 2.75 | $ 5.50 | | 1.5 | $ 2.75 | $ 4.13 | 8 transects for subset monitoring |
| 6.1 | Outplanting | Push mounts | Consumable | 200 | $ 0.07 | $ 13.48 | | 8500 | $ 0.07 | $ 572.90 | |
| 6.3 | Outplanting | Push mounts | Consumable | 200 | $ 0.07 | $ 13.48 | | 6900 | $ 0.07 | $ 465.06 | |
| 6.6 | Outplanting | Push mounts | Consumable | 200 | $ 0.07 | $ 13.48 | | 5700 | $ 0.07 | $ 384.18 | |
| 7 | Yearly monitoring | Dive gear | Equipment | 2 | $ 1,000.00 | $ 1,142.86 | Pro rata over 14 days | 2 | $ 1,000.00 | $ 444.44 | Pro rata over 18 & 11 days |
| 7 | Yearly monitoring | Slate | Equipment | 2 | $ 2.66 | $ 5.32 | | 6 | $ 2.66 | $ 15.96 | |
| 7 | Yearly monitoring | Caliper | Equipment | 2 | $ 3.50 | $ 7.00 | | 6 | $ 3.50 | $ 21.00 | |
| 7 | Yearly monitoring | Caribiner RVS | Equipment | 2 | $ 1.50 | $ 3.00 | | 6 | $ 1.50 | $ 9.00 | |
| 7 | Yearly monitoring | Pencils | Consumable | 8 | $ 0.33 | $ 2.64 | | 24 | $ 0.33 | $ 7.92 | |
| 7 | Yearly monitoring | Rubber bands wide | Consumable | 8 | $ 0.10 | $ 0.80 | | 24 | $ 0.10 | $ 2.40 | |
| 7 | Yearly monitoring | Underwater paper | Consumable | 40 | $ 1.00 | $ 39.84 | | 40 | $ 1.00 | $ 39.84 | |
| 7 | Yearly monitoring | Cattle tags maintenance | Consumable | 20 | $ 0.26 | $ 5.20 | | 20 | $ 0.26 | $ 5.20 | |
| 7 | Yearly monitoring | Masonry nails 1-1/2" (pounds) | Consumable | 4 | $ 2.75 | $ 11.00 | | 4 | $ 2.75 | $ 11.00 | |

#### Table S4. Labour & Boats

| Act | Activity | Task | Item | Personelle | Total experiment | Total scaled | Notes |
|---|---|---|---|---|---|---|---|
| 1 | Collection adult colonies | Set up holding tank | Person hours | Research assistant | 4 | 4 | |
| 1 | Collection adult colonies | Set up holding tank | Person hours | Scientific advisor | 4 | 4 | |
| 1 | Collection adult colonies | Coral collection | Person hours | Research assistant | 4 | 4 | |
| 1 | Collection adult colonies | Coral collection | Person hours | Scientific advisor | 4 | 4 | |
| 1 | Collection adult colonies | Coral collection | Boat rental & Operator half day | | 1 | 1 | |
| 1 | Collection adult colonies | Coral collection | Fuel | | 1 | 1 | |
| 1 | Collection adult colonies | Coral collection | Scuba tanks | | 4 | 4 | |
| 1 | Collection adult colonies | Coral replant | Boat rental & Operator half day | | 1 | 1 | |
| 1 | Collection adult colonies | Coral replant | Fuel | | 1 | 1 | |
| 1 | Collection adult colonies | Coral replant | Scuba tanks | | 4 | 4 | |
| 2 | Spawning & larvae rearing | Set up water filtration systems | Person hours | Research assistant | 2 | 2 | |
| 2 | Spawning & larvae rearing | Set up water filtration systems | Person hours | Scientific advisor | 2 | 2 | |
| 2 | Spawning & larvae rearing | Set up spawning materials | Person hours | Research assistant | 8 | 8 | |
| 2 | Spawning & larvae rearing | Set up spawning materials | Person hours | Scientific advisor | 8 | 8 | |
| 2 | Spawning & larvae rearing | Monitor for spawning | Person hours | Research assistant | 21 | 21 | |
| 2 | Spawning & larvae rearing | Monitor for spawning | Person hours | Scientific advisor | 21 | 21 | |
| 2 | Spawning & larvae rearing | Collection, washing & fertilizing gametes | Person hours | Research assistant | 5 | 5 | |
| 2 | Spawning & larvae rearing | Collection, washing & fertilizing gametes | Person hours | Scientific advisor | 5 | 5 | |
| 2 | Spawning & larvae rearing | Estimation of embryo density in 12 cultures | Person hours | Scientific advisor | 2 | 2 | |
| 2 | Spawning & larvae rearing | Twice daily waterchanges | Person hours | Research assistant | 14 | 56 | 2 people needed for scale |
| 2 | Spawning & larvae rearing | Set up air systems | Person hours | Research assistant | 2 | 8 | 2 people needed for scale |
| 2 | Spawning & larvae rearing | Make banjo filters | Person hours | Research assistant | 4 | 8 | |
| 2 | Spawning & larvae rearing | Adjust water flow | Person hours | Research assistant | 2 | 8 | 2 people needed for scale |
| 2 | Spawning & larvae rearing | Install airflow | Person hours | Research assistant | 2 | 8 | 2 people needed for scale |
| 3 | Substrates & Conditioning | Collect CCA | Person hours | Research assistant | 4 | 4 | |
| 3 | Substrates & Conditioning | Collect CCA | Person hours | Scientific advisor | 4 | 4 | |
| 3 | Substrates & Conditioning | Organise CAPs on rods | Person hours | Research assistant | 2 | 16 | 2 people needed for scale |
| 3 | Substrates & Conditioning | Estimation of larval densities (12 cultures) | Person hours | Scientific advisor | 2 | 2 | |
| 3 | Substrates & Conditioning | Introduce CAPs to larvae | Person hours | Research assistant | 2 | 8 | 2 people needed for scale |
| 3 | Substrates & Conditioning | Brushing/trimming CCA | Person hours | Research assistant | 1.5 | 6 | |

#### Table S4. Labour & Boats

| Act | Activity | Task | Item | Personelle | Total experiment | Total scaled | Notes |
|---|---|---|---|---|---|---|---|
| 3 | Substrates & Conditioning | Collect CCA | Boat rental & Operator half day | | 1 | 1 | |
| 3 | Substrates & Conditioning | Collect CCA | Fuel | | 1 | 1 | |
| 3 | Substrates & Conditioning | Collect CCA | Scuba tanks | | 4 | 4 | |
| 4 | Flow through 1m outplant | Set up flow through system | Person hours | Research assistant | 8 | 32 | 2 people needed for scale |
| 4 | Flow through 1m outplant | Make banjo filters | Person hours | Research assistant | 4 | 16 | 2 people needed for scale |
| 4 | Flow through 1m outplant | Adjust water flow | Person hours | Research assistant | 5 | 20 | |
| 5 | Ex-situ nursery | Build table for tanks | Person hours | Research assistant | 4 | 0 | |
| 5 | Ex-situ nursery | Set up tanks & rods | Person hours | Research assistant | 8 | 8 | |
| 5 | Ex-situ nursery | Maintenance and tank cleaning (3m) | Person hours | Research assistant | 24 | 96 | |
| 5 | Ex-situ nursery | fish & coral feeding & filter change (3m) | Person hours | Research assistant | 24 | 96 | |
| 5.6 | Ex-situ nursery | Maintenance and tank cleaning (6m) | Person hours | Research assistant | 24 | 96 | |
| 5.6 | Ex-situ nursery | fish & coral feeding & filter change (6m) | Person hours | Research assistant | 24 | 96 | |
| 6.1 | Outplanting | Prepare outplant | Person hours | Research assistant | 4 | 28 | |
| 6.1 | Outplanting | Prepare outplant | Person hours | Scientific advisor | 1 | 4 | |
| 6.1 | Outplanting | Outplant corals | Person hours | Research assistant | 8 | 520 | 5 people needed for scale |
| 6.1 | Outplanting | Outplant corals | Person hours | Scientific advisor | 8 | 104 | |
| 6.1 | Outplanting | Clean up materials | Person hours | Research assistant | 2 | 15 | |
| 6.1 | Outplanting | Outplant corals | Boat rental & Operator full day | | 1 | 13 | |
| 6.1 | Outplanting | Outplant corals | Fuel | | 1 | 13 | |
| 6.1 | Outplanting | Outplant corals | Scuba tanks | | 6 | 234 | |
| 6.3 | Outplanting | Prepare outplant | Person hours | Research assistant | 4 | 24 | |
| 6.3 | Outplanting | Prepare outplant | Person hours | Scientific advisor | 1 | 4 | |
| 6.3 | Outplanting | Outplant corals | Person hours | Research assistant | 8 | 440 | 5 people needed for scale |
| 6.3 | Outplanting | Outplant corals | Person hours | Scientific advisor | 8 | 88 | |
| 6.3 | Outplanting | Clean up materials | Person hours | Research assistant | 2 | 13 | |
| 6.3 | Outplanting | Outplant corals | Boat rental & Operator full day | | 1 | 11 | |
| 6.3 | Outplanting | Outplant corals | Fuel | | 1 | 11 | |
| 6.3 | Outplanting | Outplant corals | Scuba tanks | | 6 | 198 | |
| 6.6 | Outplanting | Prepare outplant | Person hours | Research assistant | 4 | 20 | |
| 6.6 | Outplanting | Prepare outplant | Person hours | Scientific advisor | 1 | 4 | |

#### Table S4. Labour & Boats

| Act | Activity | Task | Item | Personelle | Total experiment | Total scaled | Notes |
|---|---|---|---|---|---|---|---|
| 6.6 | Outplanting | Outplant corals | Person hours | Research assistant | 8 | 360 | 5 people needed for scale |
| 6.6 | Outplanting | Outplant corals | Person hours | Scientific advisor | 8 | 72 | |
| 6.6 | Outplanting | Clean up materials | Person hours | Research assistant | 2 | 11 | |
| 6.6 | Outplanting | Outplant corals | Boat rental & Operator full day | | 1 | 9 | |
| 6.6 | Outplanting | Outplant corals | Fuel | | 1 | 9 | |
| 6.6 | Outplanting | Outplant corals | Scuba tanks | | 6 | 162 | |
| 7 | Yearly monitoring | Monitor corals and maintain transects | Person hours | Research assistant | 48 | 32 | Monitoring subset |
| 7 | Yearly monitoring | Monitor corals and maintain transects | Person hours | Scientific advisor | 48 | 32 | Monitoring subset |
| 7 | Yearly monitoring | Data compilation | Person hours | Scientific advisor | 32 | 32 | Monitoring subset |
| 7 | Yearly monitoring | Monitor corals and maintain transects | Boat rental & Operator full day | | 4 | 4 | Monitoring subset |
| 7 | Yearly monitoring | Monitor corals and maintain transects | Boat rental & Operator half day | | 4 | 0 | Monitoring subset |
| 7 | Yearly monitoring | Monitor corals and maintain transects | Fuel | | 8 | 4 | Monitoring subset |
| 7 | Yearly monitoring | Monitor corals and maintain transects | Scuba tanks | | 40 | 24 | Monitoring subset |
